# Complex genetic patterns and distribution limits mediated by native congeners of the worldwide invasive red-eared slider turtle

**DOI:** 10.1101/2021.09.02.458785

**Authors:** Sayra Espindola, Ella Vázquez-Domínguez, Miguel Nakamura, Luis Osorio-Olvera, Enrique Martínez-Meyer, Edward A. Myers, Isaac Overcast, Brendan N. Reid, Frank T. Burbrink

**Affiliations:** Departamento de Ecología de la Biodiversidad, Instituto de Ecología, Universidad Nacional Autónoma de México, Coyoacán, Ciudad de México, 04510, México; Posgrado en Ciencias Biológicas, Universidad Nacional Autónoma de México, Coyoacán, Ciudad de México, 04510, México; Centro de Investigación en Matemáticas (CIMAT), Calle Jalisco S/N, Colonia Valenciana, 36023, Guanajuato, Guanajuato, México; Instituto de Biología, Universidad Nacional Autónoma de México, Coyoacán, Ciudad de México, 04510, México; Department of Biological Sciences, Clemson University, Clemson, SC 29634, USA; Institut de Biologie de l’Ecole Normale Superieure, 75005 Paris, France; Rutgers University, Department of Ecology, Evolution, and Natural Resources, 14 College Farm Road, New Brunswick, NJ, USA; American Museum of Natural History, Central Park West, 79th Street, New York 10024, USA

**Keywords:** niche-centrality, population genomics, species distribution models, species invasions, *Trachemys*

## Abstract

Non-native (invasive) species offer a unique opportunity to study the geographic distribution and range limits of species, wherein the evolutionary change driven by interspecific interactions between native and non-native closely related species is a key component. The red-eared slider turtle, *Trachemys scripta elegans* (TSE), has been introduced and successfully established worldwide. It can coexist with its native congeners *T. cataspila, T. venusta* and *T. taylori* in Mexico. We performed comprehensive fieldwork, executed a battery of genetic analyses and applied a novel species distribution modeling approach to evaluate their historical lineage relationships and contemporary population genetic patterns. Our findings support the historical common ancestry between native TSE and non-native (TSE_alien_), while also highlighting the genetic differentiation of the exotic lineage. Genetic patterns are associated with their range size/endemism gradient, the microendemic *T. taylori* showed significant reduced genetic diversity and high differentiation, whereas TSE_alien_ showed the highest diversity and signals of population size expansion. Counter to our expectations, lower naturally occurring distribution overlap and little admixture patterns were found between and its congeners, exhibiting reduced gene flow and clear genetic separation across neighboring species despite having zones of contact. We demonstrate that these native *Trachemys* species have distinct climatic niche suitability, likely preventing establishment of and displacement by the non-native TSE. Additionally, we found major niche overlap between TSE_alien_ and native species worldwide, supporting our prediction that sites with closer ecological optima to the invasive species have higher establishment risk than those that are closer to the niche-center of the native species.

## Introduction

How the combined effects of evolutionary, biogeographical, and ecological forces have driven the distribution of natural species, coupled with how interacting factors (e.g. physical and climatic environment, species interactions, resources availability) define the limits of species’ geographic ranges, remains a fundamental question in ecology and evolution. Non-native (invasive) species offer a unique opportunity to explore patterns and processes governing the geographic distribution and range limits of species (Goldberg and Lande, 2007; Grayson and Johnson, 2018; Espindola et al., 2019). For instance, Tingley et al. (2014) demonstrated that the rapid and extreme success of the cane toad (*Rhinella marina*) invasion of Australia is related to a shift in the species’ realized niche, as opposed to evolutionary shifts in the range-limiting traits that define its fundamental niche. An assessment of terrestrial gastropods found that native ranges reflect natural dispersal limitation and biogeographic realms, while their distribution in non-native regions is mostly explained by prevailing climate (Capinha et al., 2015).

Reconstructing human-mediated species invasion histories has been possible with the use of genetic approaches that isolate ecological and evolutionary forces acting on non-native species during the invasion process and shaping their distribution limits (Cristescu, 2015; Suárez-Atilano et al., 2019). Likewise, genetic diversity within populations of non-native species depends on a variety of factors, including the frequency of bottlenecks and founder effects, the possibility and frequency of multiple introductions and admixture, and demographic characteristics (Brennan et al., 2014; Suárez-Atilano et al., 2019). Native species also experience novel pressures when a potential invader arrives, including increased competition, range displacement and decreases in population size, introduction of maladapted alleles and potential decrease in fitness by hybridization and introgression, and even alteration of their evolutionary trajectory in response to selective pressures and rapid adaptive changes (Suarez and Tsutsui, 2008; Stuart et al., 2014; Vyšniauskienė et al., 2018). Notably, these processes have rarely been evaluated along the distribution of non-native species with their congeners, or in scenarios where they co-occur. An example illustrating how interspecific interactions between native and non-native closely related species can drive evolutionary change is shown by the lizards *Anolis carolinensis* and *A. sagrei* on islands in Florida, where the native *A. carolinensis* moved to high perches and adaptively evolved larger toepads after the *A. sagrei* invasion (Stuart et al., 2014).

Species invasions have been studied under different approaches; among these, species distribution models (SDM) have been used to assess probable risk areas based on the species’ niche and its climatic suitability (e.g. Srivastava et al., 2019). A key concept in SDM is that fitness is expected to be highest in areas with environments closest to the center of the fundamental niche (Maguire, 1973; Martínez-Meyer et al., 2013; Osorio-Olvera et al., 2019), which is defined as the physiological range of tolerance to environmental factors where a species can survive in the absence of biotic interactions (Hutchinson, 1957; Soberón and Arroyo-Peña, 2017; Espindola et al., 2019). In fact, a significant relationship between species’ fitness attributes (e.g. genetic diversity and abundance) with the distance to the center of the niche has been documented in a diverse array of taxa (Lira-Noriega and Manthey, 2014; Osorio-Olvera et al., 2020a). Contrasting niche-center distances between native and invasive species can be a useful approach to estimate the potential for establishment of the latter in sites that are suitable for both species. This idea is based on the hypothesis that the invading species would displace the native one in those sites that are closer to its niche center.

Among vertebrates, turtles represent an exceptional case for exploring genetic and distributional invasion processes, where studies have revealed complex patterns of genetic variation regarding both historical lineage diversification (Fritz et al., 2012; Kraus, 2015) and contemporary changes resulting directly from human-mediated invasions (Fong and Chen, 2010; Parham et al., 2013, 2020). The slider turtle genus *Trachemys* is one of the most widely distributed American reptile groups (Siedel and Ernst, 2017). The North American red-eared slider turtle, *Trachemys scripta elegans*, is well-known globally mainly because it has been introduced and successfully established worldwide, in at least 70 countries (CABI, 2019), due to its popularity as a reptile pet (GISD 2018). It is classified as one of the 100 worst invasive species in the World (Lowe et al., 2000), displaying an extraordinary potential for impacting native species and habitats (Cadi et al., 2004; Ficetola et al., 2009; Parham et al., 2013, 2020; Pearson et al., 2015).

*Trachemys scripta elegans* is distributed natively throughout the northeastern coast of the United States, to northern Florida, through the Mississippi River valley, west through Texas, Oklahoma, and Kansas, finally reaching its southernmost limit at northern Tamaulipas, Mexico. Other congeneric species distributed in Mexico have an interesting natural distribution that follows that of *T*.*s. elegans* in a southwards gradient along Mexico’s east coast, including native populations of *T. cataspila* (northern Tamaulipas to northern Veracruz) and *T. venusta* (northern Veracruz southwards to Guatemala. Belize and Honduras; Ernst and Siedel, 2006; Siedel and Ernst, 2017). *Trachemys taylori*, a microendemic species with a markedly restricted distribution, inhabits the wetlands system within the Cuatrociénegas desert valley, in the Chihuahua desert, Coahuila, México (Siedel, 2002a; Lazcano et al., 2019). Along this natural gradient, *T*.*s. elegans* has been introduced in different areas where it can coexist with its native congeners. The physiology and patterns of invasion of *T*.*s. elegans* have been well-studied (see Willmore and Storey, 1997; Cadi et al., 2004; Ryan et al., 2014), and with regard to phylogeny, ecology, and niche characterization (e.g. Tucker, 2001; Fritz et al., 2012; Jo et al., 2017; Espindola et al., 2019). However, historical and contemporary patterns of genetic diversity and structure remain little known in this invasive taxon, particularly comparing native and non-native populations within *T*.*s. elegans*, and among closely related species co-occurring along their distribution (but see Parham et al., 2013, 2020; Vamberger et al., 2020).

Hence, we aimed to evaluate the historical lineage relationships and contemporary population genetic patterns of these four *Trachemys* species. We specifically targeted the following questions: 1) What are the phylogenetic relationships among native and non-native *T*.*s. elegans* and in relation with its *Trachemys* congeners?, 2) Do the genetic diversity and structure patterns within and among the four species reflect their range size/endemism gradient from microendemic (*T. taylori*) to widespread invasive (*T*.*s. elegans*)?, 3) Do native turtles show signals of genetic admixture and introgression among them and with the invasive *T*.*s. elegans* in Mexico? 4) Do these *Trachemys* species exhibit distinct climatic niche suitability, enough to prevent the establishment of and displacement by the non-native *T*.*s. elegans*? and 5) Is the establishment of invasive *T*.*s. elegans* and the displacement of native turtles worldwide associated with the distance from their niches’ center?

We performed comprehensive fieldwork in Mexico and executed a battery of genetic and genomic analyses and a novel species distribution modeling approach to address these questions. We used three sets of molecular markers, microsatellite loci, mitochondrial and nuclear sequences, and single nucleotide polymorphisms (SNPs). Considering the historical dispersal and vicariance events of slider turtles (Fritz et al., 2012), as well as the history of invasion of *T*.*s. elegans* (Ficetola et al., 2009; Pearson et al., 2015; Espindola et al., 2019) and evidence of hybridization in other regions where *T*.*s. elegans* has been introduced (Parham et al., 2013, 2020), we anticipated complex genetic patterns in the *Trachemys* species studied here. Namely, we predicted disjunct population structure among the four species, historical and contemporary genetic differentiation between native and non-native *T*.*s. elegans*, and reduced genetic diversity and high structure in the microendemic *T. taylori*. We also expected to find high introgression and admixture between *T*.*s. elegans* and its congeners, which will also be supported by markedly shared suitability of their ecological niches. Finally, given that *T*.*s. elegans* has invaded and displaced native turtle species worldwide, we predicted that sites that are closer to *T*.*s. elegans*’ niche-center would have higher establishment risk than those that are closer to the niche-center of the native species.

## Materials and methods

### Field sampling, museum samples and traded individuals

We performed several field trips in 2014 and 2015 at 11 localities throughout the natural distribution of *Trachemys venusta* and *T. cataspila* along the Mexican east coast (Gulf of Mexico and the Caribbean) and *T. taylori* in the Cuatrociénegas valley. Notably, we did not find *T. scripta elegans* (TSE hereafter) at any of the naturally distributed populations with its congeneric species, where we expected it to occur given its invasive nature. Thus, we obtained TSE samples (outside its native range) only in targeted sites at artificial waterbodies (botanical gardens, parks, urban lakes), where it was undoubtedly transported intentionally. Turtles were trapped with baited hoop traps and hand nets, with a consecutive effort of 1 to 3 days per site. All captured individuals were measured (length and width of carapace and plastron) and weighted; tissue samples (∼3 mm^2^) were obtained from inter-digital tissue. All animals were released at the sampling site; fieldwork was carried out in strict adherence to the guidelines for work with amphibians and reptiles (Beaupre et al., 2004), and with the respective scientific collection permit from Secretaría del Medio Ambiente y Recursos Naturales (Semarnat-FAUT-0168). In order to have samples that covered most of TSE’s native distribution along western USA (Seidel, 2002b), 39 samples (muscle and skin) were obtained from museum specimens (the Herpetology Collections at the Field Museum of Natural History, the American Museum of Natural History, and the Smithsonian National Museum of Natural History) (Fig. 1a; Table S1). Finally, in order to gather data about traded turtles of unknown origin, 23 samples were obtained from the Herpetology Laboratory “Vivario” (FES-Iztacala, UNAM). Both fresh and museum tissue samples were stored at -20°C in 90% ethanol until later use.

**Figure 1.**
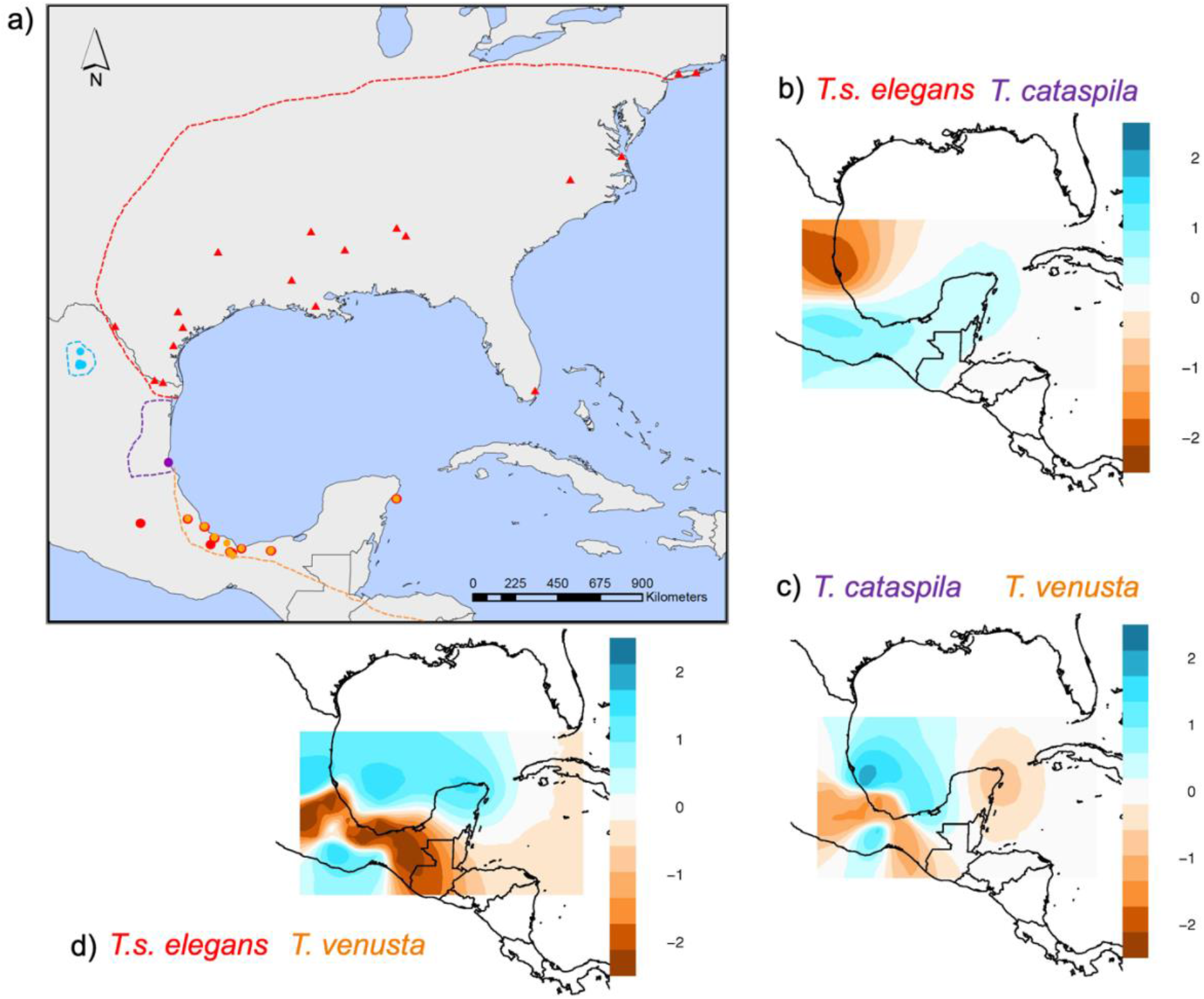
**(a)** Map of sample localities (from field work, museum collections, and the Herpetology Laboratory; Table S1) for four *Trachemys* species: *T*.*s. elegans* in its native (museum samples; red triangles) and non-native (red dots) distribution, *T. cataspila* (purple), *T. venusta* (orange) and *T. taylori* (blue). Dashed lines depict the approximate distribution range for each species. (**b-d**) Historical migration patterns (EEMS results) depicting regions of significantly reduced gene flow between the neighboring species, *T*.*s. elegans* and *T. cataspila* (**b**) and *T. cataspila* and *T. venusta* (**c**), and between the non-neighboring *T*.*s. elegans* and *T. venusta* (**d**).

### DNA extraction

We extracted total DNA from fresh tissue with the DNeasy Blood and Tissue Extraction Kit (QIAGEN), following the manufacturer’s protocols. For tissue museum samples, many of which were preserved in formaldehyde and later stored in ethanol, DNA was isolated using a modification of the QIAamp DNA FFPE Tissue Kit (QIAGEN). Before DNA extraction, samples were gradually dehydrated with ethanol, followed by a rehydration in an alkali buffer and incubation at high temperature, to eliminate formaldehyde and recover DNA fragments (Fang et al., 2002; Campos and Gilbert, 2012). DNA extractions were quantified with the Qubit fluorometer and quality was assessed with 1% agarose gels stained with GelRed (Biotium) and visualized with UV light.

### Amplification of microsatellite loci and mitochondrial and nuclear genes

Microsatellite loci were characterized with 15 fluorescently labeled primers developed for TSE (Xin et al., 2012; Simison et al., 2013). List of primers used and PCR amplification protocol are available in Table S2 and Appendix S1. Microsatellites products were run on an ABI Prism3730xl and 3100 Genetic Analyzer (Applied Biosystems), with ROX-500 as internal size standard. Allele size was determined with GeneMarker 1.97 (SoftGenetics) and rounded to integers using TANDEM 1.8 (Matschiner and Salzburger, 2009). We included negative controls in all runs and genotyped two or more randomly chosen samples to ensure correct readings and data consistency. Potential null alleles and scoring errors were assessed with MicroChecker 2.2.3 (Van Oosterhout et al., 2004), using 1,000 Monte Carlo simulations.

We also amplified the mitochondrial cytochrome *b* (cyt *b*) gene and the nuclear genomic intron 1 of the RNA fingerprint protein 35 (R35) gene for a subset of fresh tissue samples from fieldwork (no museum sample could be successfully amplified), using the same primers as in Fritz et al. (2012) and Fujita et al. (2004), respectively (Appendix S1 for PCR protocols). PCR products were sent to Macrogen-Korea for both forward and reverse sequencing.

### Genotyping by Sequencing (GBS) and SNPs calling

A subset of 178 samples was sent to the Genomic Diversity Facility (GDF) at Cornell University for Genotyping by Sequencing library construction (GBS; Elshire et al., 2011) and sequencing services. Libraries were prepared using the EcoT221 enzyme for genome complexity reduction, with recognition sequence 5’-ATGCA/T-3’ and 3’-T/ACGTA-5’. The resulting fragments were tagged with individual barcodes, samples were pooled and PCR-amplified, multiplexed, purified and sequenced on a single lane on an Illumina Hi-Seq 2500 (Illumina Inc, USA) with single-end 100 bp reads. Raw data were processed with *ipyrad*, a simple and reproducible RADseq assembly and analysis framework that is computationally efficient and suitable for both population genetic and phylogenetic scale datasets. The assembly workflow in *ipyrad* is fully self-contained, capable of taking raw Illumina data and producing assembled output files without the need for pre- or post-processing by other software (Eaton and Overcast, 2020). Our workflow was based on a *de novo* assembly method and with the filtering protocol described in Appendix S1.

### Population genetics and evolutionary relationships

To assess our questions regarding evolutionary relationships and population genetics, we performed a hierarchical set of analyses, for which we distinguished between *T. scripta elegans* individuals from its native (TSE) and non-native (TSE_alien_) distribution. First, we used the mitochondrial and nuclear sequences datasets with the purpose of determining the genetic differentiation among TSE samples from its native distribution and the non-native ones from our sampling in Mexico, as well as its divergence in relation with its *Trachemys* congeners. Thus, we performed phylogenetic inferences and estimated times of divergence. Secondly, focused on a more contemporary evaluation, we estimated genetic diversity values and population genetic structure (differentiation) patterns, based on microsatellite loci, per species and for both TSE and TSE_alien_. Finally, based on the SNPs dataset, we further evaluated phylogenetic and differentiation patterns, and estimated introgression and historical migration.

### Mitochondrial and nuclear phylogenetic analyses and divergence time estimation

For the phylogenetic analysis among the four *Trachemys* species, in addition to the sequences we amplified, we incorporated from GenBank five R35 and eight cyt *b* sequences in order to have a more complete dataset for TSE along its native distribution, plus four cyt *b* sequences for *T. scripta scripta*, one *T. venusta* and one *T. cataspila* (Table S3); we also included sequences for *Chrysemys picta, Malaclemys terrapin* and *Trachemys decussata* as outgroups, the former for both genes (cyt *b* and R35) while the other two were available only for cyt *b* (GenBank; Table S3). Sequence datasets were assembled and reviewed by eye with Bioedit 7.0.5.2 (Hall, 1999), while Geneious v.9.1.8 was used for alignment, blasting and edition.

We chose the best nucleotide substitution model for both cyt *b* and R35 datasets based on the corrected Akaike Information Criterion (AIC) in PhyML 3.0 (Guindon et al., 2010), using the default options and optimized NNI search. Phylogenetic relationships were estimated with Maximum likelihood (ML) for both genes with PhyML, using NNI+SPR search for branch length optimization and clade support was assessed with 1,000 bootstrap replicates. Posterior probabilities and a 50% majority-rule consensus trees were obtained and the resulting trees visualized with FigTree v1.4.2 (Rambaut, 2014). To evaluate the relationship at the level of mitochondrial haplotypes specifically within *T. scripta elegans/T. scripta scripta*, we constructed an unrooted network among unique cyt *b* haplotypes with PopArt (Leigh and Bryant, 2015), based on the minimum-spanning method (Bandelt et al., 1999).

Lineage divergence was estimated with the cyt *b* dataset and a Bayesian framework in Beast 1.8.3 (Suchard et al., 2018). We used the GTR model across all gene and codon positions, with empirical base frequencies and four gamma categories. We used the same outgroup species as in the phylogenetic analysis, based on which we provided calibration points and error estimates derived from a lognormal distribution, as follows: split of *Chrysemys picta* and the ingroup at 22.5 million years ago (My), *Malaclemys terrapin* and the ingroup 13.0 My (95% HPD: 9.5-17.2), *T. decussata/T. scripta* -*T. venusta* split at 9.4 My (95% HPD: 7.9-11.2), and *T. decussata-T. scripta* at 8.6 My (95% HPD: 6.9-10.5) (Fritz et al., 2012; Spinks et al., 2016). Time to the most recent common ancestor (TMRCA) was estimated with an uncorrelated lognormal tree prior and a constant population size prior, running two MCMC chains for 10×10^7^ generations, sampling every 1,000^th^ iteration. Tracer v1.7.0 was used to monitor convergence and adequate mixing (Effective Sample Size, ESS >200; Rambaut et al., 2018) and a maximum clade credibility tree (burn-in 10%) was built with TreeAnnotator v.2.4.6 (Drummond et al., 2012) and visualized with FigTree. Additionally, we conducted a skyline plot analysis using Beast to infer potential population fluctuations specifically for the non-native *T*.*s. elegans*. Model parameters were sampled each 5000 generations, with a total of 50 million generations, under a relaxed log-normal molecular clock model, with assumption of uniform distributions, root time set based on the lineage divergence results, and 10% discharged as burn-in. Convergence of results were visualized with Tracer.

### Diversity, differentiation, and demographic analyses (microsatellites loci)

We tested for deviations from Hardy-Weinberg equilibrium (HWE) and linkage disequilibrium (LD) between all pairs of loci using Genepop v.6 (Rousset, 2008) per species, with an exact probability test (Markov chain parameters: 10,000 dememorizations, 100 batches, 10,000 iterations per batch), and sequential Bonferroni corrections. In cases of deviation from HWE, we also tested whether this was attributed to heterozygosity deficit or excess. We estimated the following genetic diversity indices: observed number of alleles (*n*_*o*_), observed (*H*_*O*_) and expected (*H*_*E*_) heterozygosity, and Nei’s unbiased expected heterozygosity (*H*_*NEI*_; Nei, 1978), using Genalex v.6 (Peakall and Smouse, 2012). Allelic richness (*A*_*r*_), a rarefied measure that considers the different sampling numbers based on a standard sample size of the smallest sample with complete genotypes at all loci, was estimated with *adegenet* in R (Jombart, 2008).

We inferred population structure between and within species with two methods: a model-free multivariate method, Discriminant Analysis of Principal Components (DAPC) with *adegenet*; we also used Structure v.2.3.4 (Pritchard et al., 2000), a Bayesian approach to infer the most likely number of genetic clusters (*K*). Based on the complete database (four *Trachemys* species), we tested for *K*=1-8 genetically distinct clusters with 20 replicates for each *K*, with 500,000 iterations and a burn-in of 100,000, including *a priori* locality data, and an admixture model with correlated allele frequencies. We further examined structure within some of the identified clusters at the species level, performing new runs using the same parameters described. For all tests, the optimal *K* was calculated according to Evanno et al. (2005) with the StructureHarvester online web server (Earl and Von Holdt, 2012), while Distruct 1.1 (Rosenberg, 2004) was used to graphically display the genetic structure results.

In order to evaluate genetic differentiation between species and genetic clusters (identified with Structure; see Results), we estimated genetic divergence between populations (*F*_*ST*_) using FSTAT (Goudet, 1995) and 10,000 randomizations. Also, Nei’s genetic distance (*D*_*NEI*_) was estimated with Genalex.

### Phylogenetic relationships with SNPs

We obtained an unrooted species tree topology with a coalescent method using SVDquartets (Chifman and Kubatko, 2014), a program designed for SNP data that computes a score based on singular value decomposition of a matrix of site pattern frequencies, corresponding to a split on a phylogenetic tree. These quartet scores are used to select the best-supported topology for quartets of taxa, which in turn can be used to infer the species phylogeny using quartet methods. We ran the program with the final dataset using 160 individuals and with the default parameters. To obtain a phylogenetic tree incorporating an outgroup, we built a SNP-based outgroup from the Painted turtle *Chrysemys picta* genome (Badenhorst et al., 2015) with Samtools 1.6 (Li, 2011). Next, we used FastTree (Price et al., 2009) to infer an approximately-maximum-likelihood phylogenetic tree, using the GTR model and the gamma option to rescale the branch lengths.

### Individual ancestry, migration, and potential introgression (SNPs)

With the aim of evaluating genetic structure at a fine scale based on SNPs, we performed a DAPC analysis with *adegenet* and secondly used the sparse non-negative matrix factorization method (sNMF; Frichot et al., 2014), which computes regularized least-square estimates of admixture proportion to estimate individual ancestry coefficients. sNMF advantages include computing homozygous and heterozygous frequencies without Hardy-Weinberg equilibrium assumptions, and adequate performance with presence of inbreeding, low sample size, large datasets and missing data (Frichot et al., 2014; Wollstein and Lao, 2015). We ran sNMF with the command-line version, *K* from 2 to 7, 6 series, and default parameters. The optimum *K* was calculated by minimal cross-entropy and results plotted as a barplot. Individuals are considered admixed if they show >75% ancestry.

The historical migration patterns were evaluated with the method of Estimated Effective Migration Surfaces (EEMS; Petkova et al., 2016). Migration patterns are adjusted by the similarity between the observed genetic differences and those expected under a stepping-stone model; the obtained estimations are interpolated across a uniformly spaced grid (the landscape) to create the migration surfaces. Genetic differentiation is represented as a function of migration rates, correlating genetic variation with geography; portions of a species range where population divergence deviates from patterns expected under isolation-by-distance are highlighted. If genetic similarity decreases rapidly for the observed, in comparison with expected values, indicates a low effective migration (i.e. a barrier to gene flow; Petkova et al., 2016). We built a pairwise dissimilarity matrix and a geographic distance matrix for all individuals to perform paired comparisons (TSE-TC; TC-TV; TSE-TV). With these matrices we ran EEMS with runeems_snps, using a deme size of 1200, with three independent starting chains for 5×10^6^ MCMC iterations, 5×10^4^ of burn-in, thinning of 5000 and different starting seeds. The three runs were combined and visualized with reemsplots2 in R (https://github.com/dipetkov/reemsplots2).

To test for introgression among species we used the abba/baba test (also called D statistics), which is a powerful test for a deviation from a strict bifurcating evolutionary history, namely testing for an excess of shared derived variants between non-sister taxa (Durand et al., 2011; Zheng and Janke, 2018). In brief, consider three ingroup populations (or species) and an outgroup with the relationship (((P1,P2),P3),O), where P1 and P2 are more related to one another than either to P3; given a haploid genome sequence for each population (i.e. H1, H2 and H3), abba sites are those at which H2 and H3 share a derived allele ‘B’, whereas H1 has the ancestral state ‘A’, as defined by the outgroup sample. Similarly, baba represents sites at which H1 and H3 share the derived state. We built three hypotheses [(TC,TV),TSE; (TT,TC),TSE; (TV,TT),TSE) plus a null model, using two randomly chosen individuals per species (to test each hypothesis twice); *Chrysemys picta* was used as outgroup. To run abba/baba we used *evobiR* in R (http://www.uta.edu/karyodb/evobiR/) with the CalcD function and a jackknife (1000) significance test.

### Niche overlap and niche suitability between TSE and native species

To evaluate our last questions about the establishment of invasive *T*.*s. elegans* and the displacement of native turtles, we explored niche suitability and niche overlap between TSE and its congeners *T. venusta, T. cataspila* and *T. taylori*; while for worldwide comparisons, we selected a set of native turtle species from different parts of the world where invasive TSE has been documented and negative interactions have been observed: *Actinemys marmorata* (Western United States; Spinks et al., 2003; Silbernagel et al., 2013), *Mauremys leprosa* (south Europe-north Africa; Polo-Cavia et al., 2014; Meyer et al., 2015), *Emys orbicularis* (Europe; Cady and Joly, 2004; Polo-Cavia et al., 2014*), Mauremys reevesii* (Asia; Jo et al., 2017; Geng et al., 2018), *Emydura macquarii* and *Chelodina longicollis* (Australia; Burgin, 2006; Mo, 2019). A dataset of available occurrence records per species was gathered from the Global Biodiversity Information Facility (GBIF, 2015); our own collecting records in Mexico were also included. To clean this dataset, we removed fossil records, doubtful occurrences, wrongly georeferenced localities and duplicate records. Then, we thinned the occurrences to avoid spatial autocorrelation problems, using a distance of 5 km, based on an average movement distance. The final dataset included the following records per species: *T*.*s. elegans n* = 4270, *T. venusta n* = 104, *T. cataspila n* = 26, *T. taylori n* = 32, *A. marmorata n* = 648, *E. orbicularis n* = 752, *M. leprosa n* = 1535, *M. reevesii n* = 219, *E. macquarii n* = 642, and *C. longicollis n* = 1559.

The environmental data used to model niches and estimate the proportion of overlap was derived from a principal component analysis on the 19 bioclimatic variables from WorldClim v1.4, at 2.5’ spatial resolution (∼5 km^2^ cell; Hijmans et al., 2005). We adopted the first three principal components (PCs), which summarized ∼85% of the total variance of the original WorldClim dataset (Fig. S1) to describe ecological niches. Niches were determined in environmental space by means of ellipsoid models based on the three-dimensional normal distribution (Osorio-Olvera et al., 2020a, 2020b) applied to the first three PCs. This calls for estimating a niche center and a positive definite covariance matrix to represent the niche, based on the occurrence points of each species. The modeled niches are each 95% highest density regions (HDR) in three dimensions, which are in fact ellipsoids because of the multivariate normal assumption (see Appendices S2-Glossary and S3-HDR). Let us denote by *A* the ellipsoid corresponding to TSE having center *μ* and covariance Σ, and by *B*_*k*_ the ellipsoids of each of the other turtles, having centers *μ*_*k*_ and covariances Σ_*k*_. The R package *ntbox* (Osorio-Olvera et al., 2020b) provides functions for estimating these niche centers and covariance matrices.

Quantifications of niche overlap are the volumes of the intersections *A* ∩ *B*_*k*_, that were approximated numerically by Monte Carlo integration as follows: by first examining the range of each PC dimension (PC_*j*_, *j* = 1, 2, 3), a hyperrectangle was constructed that contained the union *A* ∪ *B*_*k*_, and a large number of points (5 × 10^6^) was uniformly generated within the hyperrectangle. The proportion of points that randomly lie in *A* ∩ *B*_*k*_ multiplied by the volume of the hyperrectangle is an approximation to the volume of *A* ∩ *B*_*k*_. We used *ntbox* for this numerical purpose, as functions are provided to easily check if a given random point is within a given ellipsoidal niche or not.

Once niche centers and covariances were determined for each species, we addressed relative appropriateness of occurrences that lie in intersections *A* ∩ *B*_*k*_ by computing suitability indexes. Consider *M*(*p, μ*, Σ), the Mahalanobis distance between point *p* and the multivariate normal distribution with mean *μ* and covariance matrix Σ. That is, if the coordinates of *p* are (*x, y, z*), *M*(*p, μ*, Σ) = [(*x, y, z*)^*T*^ − *μ*]^*T*^Σ^−1^[(*x, y, z*)^*T*^ − *μ*]. For every occurrence point *p*_*i*_ ∈ *A* ∩ *B*_*K*_ of TSE let *M*_*1i*_ = *M*(*p*_*i*_, *μ*, Σ) and *M*_*2i*_ = *M*(*p*_*i*_, *μ*_*k*_, Σ_*k*_). For every occurrence point *q*_*j*_ ∈ *A* ∩ *B*_*k*_ of species *k*, let *M*_1*j*_ = *M*(*q*_*j*_, *μ, Σ*) and *M*_2*j*_ = *M*(*q*_*j*_, *μ*_*k*_, *Σ*_*k*_). Suitability values at a given occurrence point are defined as *S* = *e*^−*M*/2^, which rescales Mahalanobis distances to (0,1), where *M* = 0 (maximum suitability) corresponds to *S* = 1, and suitability decreases the further away a point is located from the corresponding niche center. Pairs of values (*S, S*_2*i*_) and (*S*_1*j*_, *S*_2*j*_) can thus be obtained and plotted on the unit square, the suitability space. We refer to these plots as suitability maps for each species.

A main point of interest is to study the observed bivariate distributions of (*S*_1*i*_, *S*_2*i*_) and (*S*_1*j*_, *S*_2*j*_) and compare them with what these distributions would have been under the hypothesis of independence, namely no association of the species’ distributions when the invasive species is present. If TSE and species *k* are statistically independent of each other, then the multivariate normals that determine niche location for each species are independent. This in turn induces a specific but unspecified behavior or density for (*S*_1*i*_, *S*_2*i*_) and (*S*_1*j*_, *S*_2*j*_) in the intersection *A* ∩ *B*_*k*_. This density is analytically unknown but can be studied by Monte Carlo simulation.

We employed three descriptive statistics to help discern differences between the observed behavior of (*S*_1*i*_, *S*_2*i*_) and (*S*_1*j*_, *S*_2*j*_) relative to the behavior that would be expected under independence. First, if *f*(*S*_1_, *S*_2_) is the (bivariate) density of (*S*_1*i*_, *S*_2*i*_) and *f*_*k*_ (*S*_1_, *S*_2_) is the density of (*S*_1*j*_, *S*_2*j*_), we define a distance between these two densities as

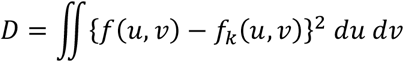

This will enable us to examine what the effect of species interaction is in terms of their relative locations and probabilities in suitability space. Secondly, we consider the sizes (areas) of the 95% HDRs for *f* and *f*_*k*_. These are denoted by *H*_1_ and *H*_2_ and are useful to examine how the interaction individually altered the way in which species distribute with high probability in suitability space. Of course, *f* and *f*_*k*_ are unknown, but since we have data observed over *A* ∩ *B*_*k*_, we can use them to compute density estimates 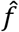 and 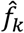 using kernel density estimation (see Appendix S1). The plots of these density estimates are useful descriptors, but in turn, they also allowed us to obtain estimates 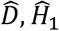, and *Ĥ*_2_ for each instance of *k*.

Statistical significance of 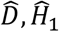, and *Ĥ*_2_ is determined by Monte Carlo simulation under the hypothesis of independence. This means to independently simulate pairs of 3-variate normal samples (of the same size as observed) within *A* ∩ *B*_*k*_, convert sampled points to suitabilities, compute kernel density estimates 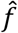 and 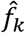, and thereby collect a large number of sample values (5000 in our study) of 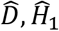, and *Ĥ*_2_. The distribution of these simulated values under independence were used to compute *p*-values by comparing to the corresponding observed values. Both upper and lower-tailed *p*-values were considered in order to determine if observed values are significant in a single-tail sense.

## Results

Our dataset included 261 samples, comprising individuals from the field, traded turtles of unknown origin, and museum samples. We could not classify five individuals to species in the field due to a markedly mixed external morphology (TSX01-TSX05; Fig. S2); these were later identified with the genomic data (see below). Thus, we worked with a final dataset of 69 *T. venusta*, 13 *T. cataspila*, 12 *T. taylori*, 128 TSE_alien_ and 39 TSE (museum) turtle individuals (Table S1).

### Sequence-based phylogeny and times of divergence

We successfully amplified 57 cyt *b* sequences (782 bp) and 41 R35 (800 bp), encompassing samples from all sampling localities in Mexico and the four *Trachemys* species studied. The best nucleotide substitution models were GTR and HKY85 for cyt *b* and R35, respectively. The ML topology with both genes consistently showed two well-differentiated clades, one for TSE (native and non-native) and the second encompassing the other three species (Fig. S3). Based on the nuclear R35 gene (Fig. S3a), TSE included eight haplotypes (2 and 4 exclusive for TSE and TSE_alien_ respectively, and 2 shared); *T. venusta* formed a separated group (4 haplotypes), while *T. cataspila* and *T. taylori* were not well resolved, each had two exclusive haplotypes while sharing one. Cyt *b* included seven haplotypes, 2 (TSE) and 3 (TSE_alien_) exclusive and 2 shared; it had low resolution for the other three species, with no clear separation among them (Fig. S3b). Results from the minimum spanning haplotype network for TSE showed one central, most abundant haplotype, which includes both native and non-native samples. A few unique terminal haplotypes, one widespread, encompass non-native haplotypes separated by one mutation step, while the others separated by >3-4 mutations are native (Illinois) or lead to *T*.*s. scripta* (Fig. S3c).

The topology obtained with Beast for the estimation of the times of divergence is concordant with the ML phylogenetic relationships depicting two distinct clades, one for TSE (native and non-native) and the other where only *T. venusta* is more clearly separated (Fig. S4). The divergence of the *Trachemys* haplotypes (node A in Fig. S4) occurred 12.9 (95% HPD: 12.6-13.3) million years ago (My). The time to the most recent common ancestor (TMRCA) leading to *T. venusta* and to *T. taylori*/*T. cataspila* was dated at 9.5 My (9.3-9.7, node B) and 9.4 My (9.2-9.7, node C), respectively; diversification times for each clade (TV and TT/TC) started ∼1.7 and ∼2.2 My respectively. The TMRCA between *T*.*s. scripta* and *T*.*s. elegans* (node D) was 8.4 My (8.1-8.6), followed by a wide range of diverging times within the TSE’s clade, which diverged at 3.2 My (node E) into two monophyletic groups, one including native and non-native haplotypes (1.9 My) and a younger one, exclusively of non-native TSE, from 1.6 My to as recent as few thousand years ago. The TSE_alien_ skyline plot showed a pattern of population size expansion towards the present (Fig. S4). Both analyses had convergence with ESS >200.

### Genetic diversity, structure, and bottlenecks (microsatellite loci)

We successfully amplified all 15 polymorphic microsatellite loci for 256 individuals. Null alleles were detected for two loci (TSE06 and TSE02) but based on the Brookfield 1 index results (<15% null alleles), all loci were retained for further analyses. After Bonferroni correction, deviations from HWE were observed at multiple loci and for each species showing in all cases significant heterozygote deficiency (*p* < 0.001). No linked loci were found. Regarding genetic variability results, allelic richness (*A*_*r*_) ranged from 3.1 (*T. taylori*) to 10.9 (TSE_alien_) (Table 1). With the exception of *T. taylori* (*H*_*O*_ = 0.255; *H*_*NEI*_ = 0.424), moderate to high observed and expected heterozygosity values were observed for all species, ranging from *H*_*O*_ = 0.580 (TSE) to *H*_*O*_ = 0.792 (TSE_alien_) and *H*_*NEI*_ = 0.767 (*T. cataspila*) and *H*_*E*_ = 0.890 (TSE_alien_). Lower genetic diversity indices in *T. taylori* were statistically significant in comparison with the other species (Mann-Whitney U tests, *p* = 0.003).

**Table 1.** Genetic diversity for *Trachemys* species analyzed in the study (five genetic clusters identified; see Structure results), based on 15 microsatellite loci. TSE and TSE_alien_: native and non-native *Trachemys scripta elegans*, respectively. Number of individual samples (N) observed (*n*_*o*_) number of alleles, allelic richness (*A*_*r*_), observed (*H*_*O*_) and expected (*H*_*E*_) heterozygosity, Nei’s unbiased expected heterozygosity (*H*_*NEI*_), and fixation index (*F*_*IS*_).

| | N | $n_o$ | $A_r$ | $H_o$ | $H_E$ | $H_{NEI}$ | $F_{IS}$ |
| --- | --- | --- | --- | --- | --- | --- | --- |
| TSE | 39 | 13.40 | 9.71 | 0.580 | 0.861 | 0.884 | 0.327 |
| TSE<sub>(alien)</sub> | 128 | 20.87 | 10.89 | 0.792 | 0.890 | 0.893 | 0.107 |
| *T. venusta* | 64 | 16.27 | 8.92 | 0.620 | 0.788 | 0.794 | 0.215 |
| *T. cataspila* | 13 | 8.02 | 7.10 | 0.626 | 0.737 | 0.767 | 0.151 |
| *T. taylori* | 12 | 3.33 | 3.15 | 0.255 | 0.406 | 0.424 | 0.331 |

Consistent results were provided by both approaches used to evaluate genetic structuring. First, the four *Trachemys* species were separated as independent genetic groups with the DAPC analysis (Fig. 2a). Similarly, Structure results clustered each species separately, although interestingly it identified five clusters (LnPr(*K* = 5) = -1780.4), clearly separating the native (TSE) and non-native individuals (TSE_alien_) as two different clusters (Fig. 2b). In a second hierarchical Structure (per species), five clusters were found within *T. venusta* (LnPr(*K* = 5) = -3571.5; Fig. 2c). In agreement, strong allelic and genetic differentiation was found (G exact test; *p* < 0.05), where the highest pairwise differentiation was found between *T. taylori* and the other species/genetic clusters (*F*_*ST*_ = 0.163 to 0.184, *D*_*NEI*_ = 0.739 to 1.012), and the lowest between TSE and TSE_alien_ with both indices (Table 2).

**Figure 2.**
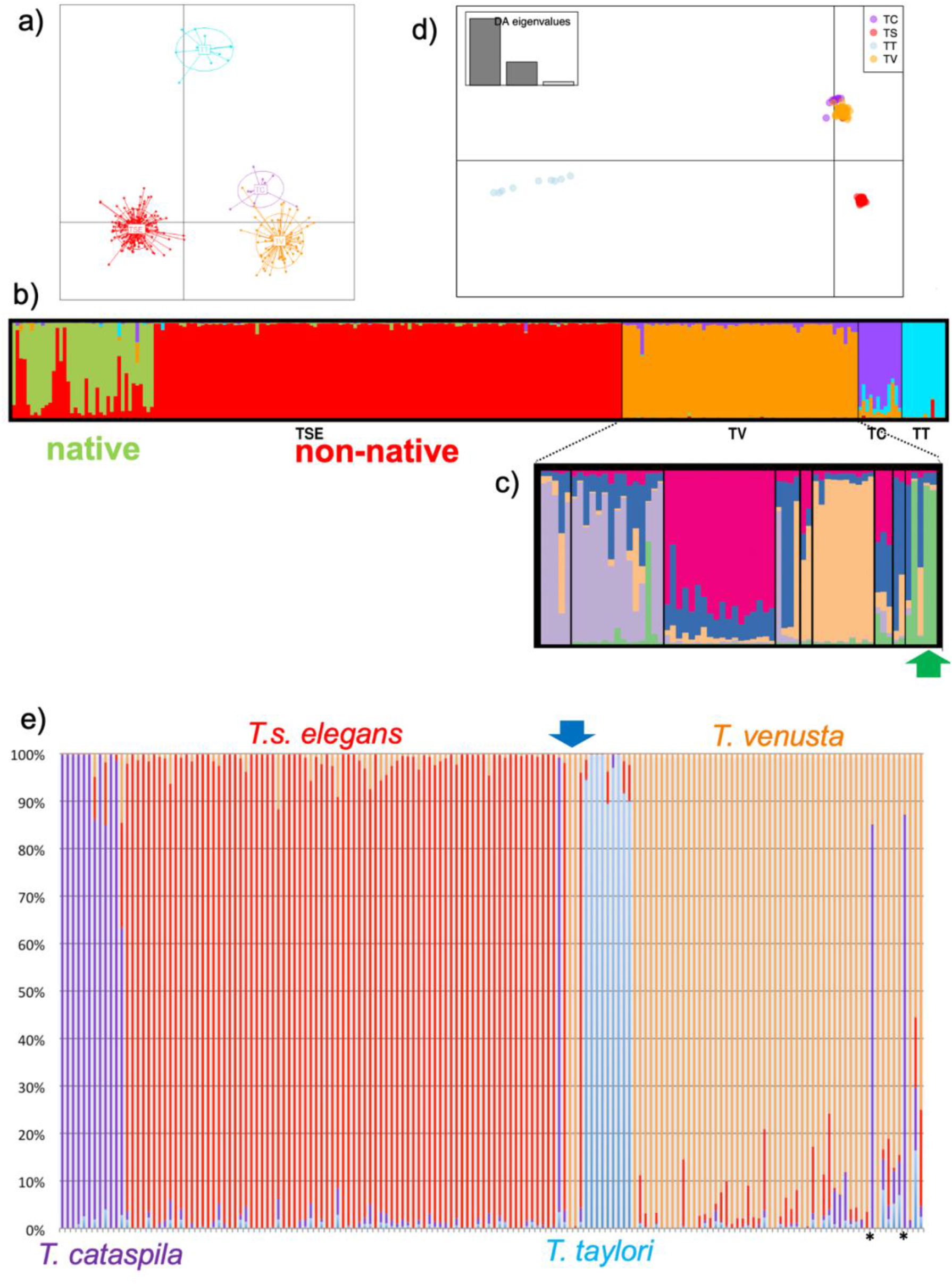
Genetic differentiation based on (**a**) DAPC analysis where the four *Trachemys* species were separated as independent genetic groups, and (**b**) Structure (*K* = 5), where the native (TSE) and non-native individuals (TSE_alien_), and also (**c**) *Trachemys venusta*, were separated into 2 and 5 different clusters, respectively (based on 15 microsatellite loci); green arrow signals individuals from Cozumel Island. (**d**) DAPC and (**e**) ancestry proportions results based on individual ancestry coefficients (sNMF) for the four *Trachemys* species (based on 63,385 SNPs). Blue arrow indicates the five individuals not identified on the field (TSX01=TC; TSX03, TSX04=TV; TSX02, TSX05=TSE) and the asterisks depict highly admixed individuals (*T. venusta* TV56 and TV64 with *T. cataspila* TC).

**Table 2.** Genetic differentiation between *Trachemys* species analyzed in the study (five genetic clusters identified; see Structure results), measured as *F*_*ST*_ (above diagonal) and Nei’s genetic distance (below diagonal), based on 15 microsatellite loci. TSE and TSE_alien_: native and non-native *Trachemys scripta elegans*, respectively.

|  | TSE | TSE<sub>(alien)</sub> | *T. venusta* | *T. cataspila* | *T. taylori* |
| --- | --- | --- | --- | --- | --- |
| TSE | - | 0.030 | 0.050 | 0.070 | 0.180 |
| TSE<sub>(alien)</sub> | 0.547 | - | 0.057 | 0.076 | 0.167 |
| *T. venusta* | 0.626 | 0.853 | - | 0.055 | 0.163 |
| *T. cataspila* | 0.692 | 0.871 | 0.395 | - | 0.184 |
| *T. taylori* | 1.012 | 0.887 | 0.739 | 0.783 | - |

### Species relationships, genomic ancestry, migration, and introgression

Results from GBS rendered a total of 483×10^6^ raw reads, an average number of raw reads per sample of 3.02×10^6^, and 210,809 raw SNPs. The final dataset after filtering included 160 samples (none for the TSE museum samples), with an average number of loci/sample = 55,982 and 63,385 SNPs. Based on this SNPs dataset, both the coalescent unrooted species (SVDquartets) and the approximately-maximum-likelihood (FastTree) phylogenetic trees (Fig. 3) exhibited a better resolved topology (in comparison with that from cyt *b* and R35 sequences), clearly separating the four species. This was further supported by the DAPC and sNMF results that showed four clusters as the optimum genetic structure (Fig. 2d,e), concordant with the four species. The SVDquartets tree showed some individuals with potential admixture (*T. venusta* TV56 and TV64 with *T. cataspila* TC), which was confirmed by sNMF where TV56 and TV64 showed high admixture with TC (>75% TC ancestry). FastTree and sNMF results also allowed us to correctly identify the five individuals we were not able to classify on the field (Fig. S2): TSX01=TC; TSX03, TSX04=TV; TSX02, TSX05=TSE. We performed an sNMF analysis for each TV and TSE separately, obtaining *K* = 3 and *K* = 2 respectively, where the former separated three TV samples from Cozumel Island (TV58, TV60, TV61) (Fig. S5a); the two TSE genetic clusters had no geographic pattern (Fig. S5b).

**Figure 3.**
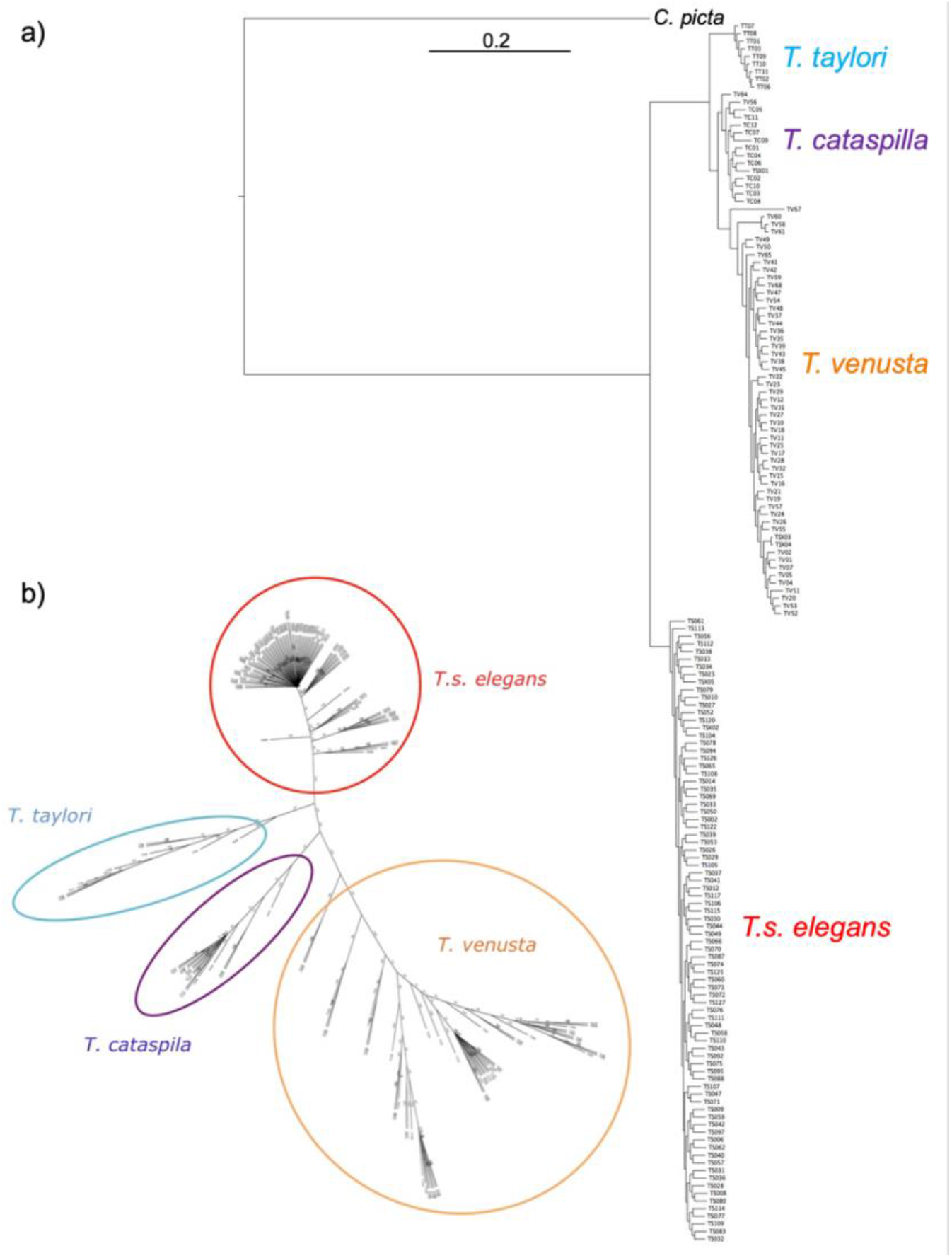
(**a**) Approximately-maximum-likelihood phylogenetic tree (FastTree) and (**b**) Unrooted species tree built with a coalescent method (SVDquartets), based on 63,385 SNPs of *Trachemys* species (*T. scripta elegans, T. cataspila, T. venusta, T. taylori*); *Chrysemys picta* as outgroup in (**a**).

Regarding historical migration patterns, the EEMS results exhibited regions of lower than expected migration across the geographic distributions of the species. Specifically, regions of significantly reduced gene flow are indicated between neighboring species, *T*.*s. elegans* and *T. cataspila* (Fig. 1b) and between *T. cataspila*-*T. venusta* (Fig. 1c); while also between *T*.*s. elegans* and *T. venusta* (Fig. 1d). Introgression results based on the *D* statistics (abba/baba test) showed signal of historical introgression between *T. cataspila* and *T*.*s. elegans* (Table S4) which share a natural contact zone, and not between other congeners.

### Niche overlap and geographic range structure

Based on niche simulations, distinct patterns were observed when overlapping climatic niches between TSE and native turtle species from different parts of the world. Results showed an overlap (respect to the native species) from 100% with *Mauremys leprosa* (i.e., the climatic niche of the native species is completely within the climatic niche of the non-native species) to 45% with *Mauremys reevesii*, which has a niche larger than TSE (Fig. S6). In geographic space, niche suitability (understood as maps depicting the proximity to the niches’ centers) showed larger suitable areas for the non-native species, except in *Mauremys reevesii*, where it was noticeably higher for the native species (*p* < 2.2e-16) (Fig. S6 [5a-5b]). When we examined the effect of species interactions, a significant change was observed both in their relative location and in the size of their HDRs (*p* < 0.003). With the exception of *Actinemys marmorata*, the change included a greater distance and a smaller area but conserving a marked overlap between the invasive and the native species (Fig. S7; Table S5).

Results for *Trachemys* species from Mexico were different. The niche overlap (respect to the native congeneric species) was 100% for *T. taylori*, 66% for *T. cataspila* and 23% for *T. venusta* (Fig. 4a). In contrast to the results obtained regarding native turtles worldwide, a larger suitable area was observed for native congeners in Mexico (*p* < 2.2e-16), coinciding with the natural distributional area occupied by each species (Fig. 4b). Additionally, a significant change was observed when bivariate distributions were compared, with a greater distance and a smaller area (*p* < 0.003), but this time the overlap zones decreased considerably (Fig. 5).

**Figure 4.**
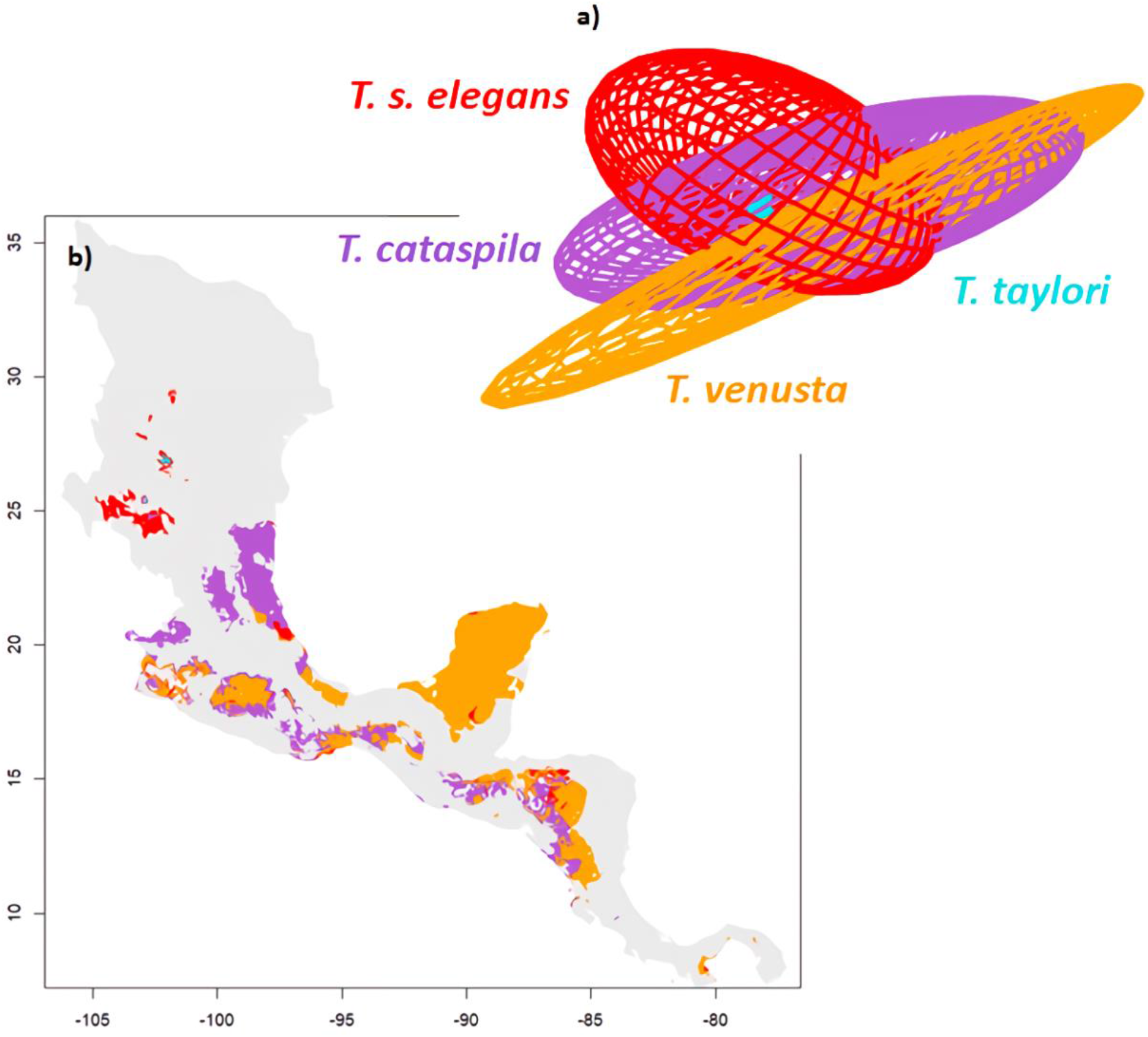
Niche overlap between the four *Trachemys* species. (**a**) Graphic representation of ecological niche overlap, based on ellipsoid models, depicting *T. scripta elegans* (red) and each native species from Mexico (*T. taylori*, blue; *T. cataspila*, purple; *T. venusta*, orange). (**b**) Map showing the sites along the distribution of each native species that are closer to the niche-center of *T. scripta elegans*. If the site has a greater suitability (understood as the proximity to the niche-center) for *T. scripta elegans* is colored red, otherwise, it is colored according to that of each native species (as in **a**). Gray color depicts areas where no overlap occurs.

**Figure 5.**
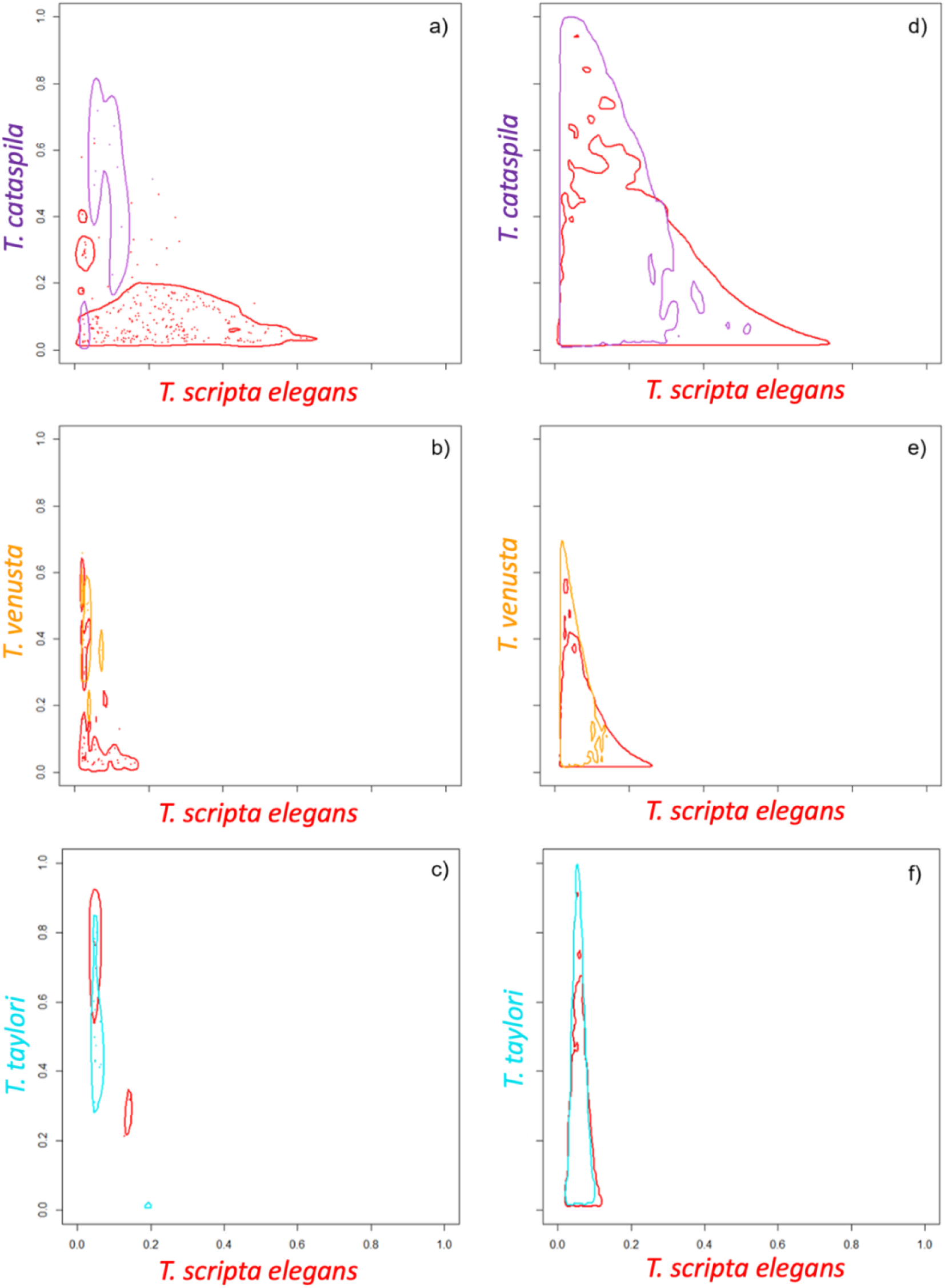
Suitability maps for *Trachemys scripta elegans* (in red) and its congeners *T. cataspila* (purple), *T. venusta* (orange) and *T. taylori* (blue). The panel on the right (**d**-**f**) shows what the highest density regions (HDRs) would be like for densities *f* and *f*_*k*_ if species are distributed independently of each other. This is obtained by Monte Carlo simulations, generating a large number of independent samples and applying kernel density estimates. By contrast, the left panel (**a**-**c**) is the suitability map based on observed data over *A* ∩ *B*_*k*_.

## Discussion

Understanding how niche differences affect biological invasions and their genetic consequences is crucial for conserving native biodiversity. We present the first assessment of the genetic and ecological niche patterns of *Trachemys script elegans*, a widely known invasive, and three of its native congeneric *Trachemys* species. As predicted, our findings showed historical and contemporary divergence between native and non-native *T*.*s. elegans* and reduced genetic diversity and high differentiation in the microendemic *T. taylori*. Notably, contrary to our expectation, they did not show shared suitability of their ecological niches neither high introgression or admixture.

### Complex genetic patterns and disjunct population structure

Our first question aimed to evaluate the evolutionary relationships among native and non-native *T*.*s. elegans* (TSE) and with lineages of its *Trachemys* congeners. Fritz et al. (2012) and Parham et al. (2013) had each assessed three of the four species we studied; the phylogenetic relationships we evaluated of these four *Trachemys* showed that TSE is a distinct lineage from the other species; notably, native (TSE) and non-native (TSE_alien_) include both exclusive and shared cyt *b* and R35 haplotypes, indicative of their historical common ancestry while also showing divergence between them. This is further supported by the more contemporary differentiation (microsatellites), where the native and non-native individuals are clearly separated as two distinct genetic clusters.

Our divergence results agree with the historical biogeography described by Fritz et al. (2012), which showed that *Trachemys* dispersed from North America to Central America (>10 My). The two species groups diverged within Central America around 8 My, generating *T. scripta* (>5.5 My), *T. venusta* and *T. cataspila*, the last two diverging much later (2-1 My). We identified two lineages within the TSE clade, one with native and no-native haplotypes and the other with exclusively non-native ones, again highlighting the genetic differentiation of the exotic lineage, which exhibits the highest diversification (different haplotypes) mostly spanning contemporary times. Moreover, the relationships among TSE haplotypes are in agreement with this pattern of more recent, unique non-native haplotypes.

We next asked if the genetic diversity and structure patterns within and among species would reflect their range size/endemism gradient from microendemic (*T. taylori*) to the widespread invasive TSE_alien_ and predicted that *T. taylori* will show reduced genetic diversity and high differentiation. Indeed, clear genetic structure of *T. taylori* is exhibited by DAPC using both microsatellite loci and SNPs which depict it as a unique genetic cluster. In fact, the four *Trachemys* species were separated as independent genetic groups. *Trachemys taylori*, microendemic to the desert-spring ecosystem of the Cuatrociénegas Basin (CCB), Coahuila, Mexico, is only found in permanent and seasonal wetlands across the CCB valley (Siedel, 2002a). The CCB wetland has shrunk 90% in the last 60 years, mainly as a result of water extraction and channeling through a complex system of artificial courses and canals, significantly reducing the natural distribution of many species, including other endemic turtles (*Terrapene coahuila, Apalone atra*; Cortés-Rodríguez et al., 2021). Very little is known about the biology and ecology of *T. taylori*, but there is no doubt that they have rather low population sizes, likely due to their endemism but also to the contemporary habitat loss and fragmentation across the CCB, as recently described for *T. coahuila* (Cortés-Rodríguez et al., 2021). Accordingly, *T. taylori* are highly diverged from other *Trachemys* species and they present the lowest genetic diversity.

In contrast, but not unexpectedly, the highest genetic diversity was shown by the TSE_alien_ individuals, jointly with a signal of population size expansion. This can be associated with diverse factors, namely that they originated historically from distinct geographic locations and/or from multiple sources or lineages, and likely also from multiple introductions (Kolbe et al., 2007; Simison et al., 2013; Suárez-Atilano et al., 2019).

### Distribution limits mediated by native congeners

We aimed to determine the invasion scenario of TSE along its distribution gradient in Mexico in relation with its congeners. We expected that they will show high genetic admixture and introgression, which will also be evidenced by markedly shared suitability of their ecological niches. Counter to our expectations, we found lower than anticipated naturally occurring distribution overlap between TSE_alien_ and its congeners along the Gulf of Mexico coastal region, a pattern supported by different results. During our extensive fieldwork, we did not find TSE_alien_ individuals at the naturally distributed areas of its congeneric species. Also, we show that the four species are clearly genetically differentiated, while non-native TSE individuals comprise a unique genetic cluster. In addition, we identified little admixture between TSE_alien_ and any of its congeners. Parham et al. (2020) found evidence of admixture between the subspecies *T. scripta elegans* and *T. scripta scripta*, and between the former and *T. gaigeae*, which in addition share morphological features. We found two *T. venusta* individuals having high admixture with *T. cataspila*; these are also recognized as subspecies, *T. venusta venusta* and *T. venusta cataspila* (Fritz et al., 2012; Parham et al., 2013), likely due to their potential to hybridize, a process that would need to be further evaluated but out of the scope of our study. Interestingly, *T. venusta* is genetically differentiated as five clusters (with the microsatellite data) that are roughly associated with their geographic distribution, from northern Veracruz to southern Tabasco and a population from the Caribbean island Cozumel as another cluster (some of the Cozumel individuals were also differentiated by SNPs; Fig. 2, Fig. S5a).

Our findings show regions of reduced gene flow precisely across neighboring species, that is between northern TSE and *T. cataspila*, and between the latter and *T. venusta*, in agreement with their natural gradient distribution southwards along the Gulf of Mexico and the Yucatan peninsula in Mexico (Fig. 1), exhibiting clear genetic separation despite having zones of contact. Gene flow is most likely ancient rather than recent, given the geographic and genetic structuring among them, with introgression occurring in the past. Indeed, we found that the most likely past introgression was between *T. cataspila* and TSE, while *T. cataspila* and *T. venusta*, which are sister species and neighbors geographically (contact zone in southern Tamaulipas), exhibit significant low migration between them.

### Suitability, environmental centrality, and establishment risk

Comparative studies between native and invasive vertebrates that explicitly include biological interactions or functional features are quite limited (Polo-Cavia et al., 2014; Espindola et al., 2019). Thus, contrasting niche suitability could be a useful tool to include, indirectly, the interaction that can occur when a species is introduced to habitats occupied by other species, based on the environmental centrality hypothesis, where fitness is expected to be highest in those sites with environments closest to the center of the fundamental niche (Martínez-Meyer et al., 2013; Osorio-Olvera et al., 2019). To further assess the observed distribution limits and low admixture patterns mediated by TSE’s congeners, we used our novel distribution modeling approach to determine if the congeneric *Trachemys* species exhibited distinct climatic niche suitability, likely preventing establishment of and displacement by the non-native TSE. Our results suggest strongly that this is the case; we demonstrate that the niche overlap between non-native TSE and four out of the six worldwide native species we analyzed is >90%. Specifically, the climatic niche of *Mauremys leprosa* and *Emys orbicularis* (native to Europe), *Actinemys marmorata* (Western coast of USA and Mexico), and *Chelodina longicollis* (Australia) are completely (or mostly) within the climatic niche of the non-native species (TSE). Thus, the available climatic niches that TSE has encountered in those regions likely facilitated its successful establishment. The latter, jointly with specific biological and life history traits, potentially enabled it to displace the native species (Cady and Joly, 2004; Meyer et al., 2015). In fact, Lambert et al. (2019) provide evidence that TSE can compete with native *Actinemys marmorata* in the wild. What we found for *Trachemys* species from Mexico is markedly different. Niche overlap was 66% with *T. cataspila* and 23% with *T. venusta*, which follows the natural geographic climatic cline along their distribution. Surprisingly, niche overlap with *T. taylori* was 100%, despite the rather unique ecosystem of the Cuatrociénegas wetlands it inhabits. This is a crucial result, because it shows that non-native TSE, at least climatically, could easily adapt to the CCB habitat, and become a threat to the micro-endemic *T. taylori*. During our fieldwork we did not find TSE individuals, but it has been nonetheless documented (McGaugh, 2012).

Because TSE has invaded and displaced native turtle species worldwide, we predicted that sites that are closer to its niche-center would have higher establishment risk than those that are closer to the niche-center of the native species. Our findings fully support our prediction; niche suitability, understood as the proximity to the center of the niche, was greater for the non-native species in all our comparisons with exception of *Mauremys reevesii* (native to Asia), which also showed the lowest niche overlap. This turtle is known for its ability to hybridize with other Geoemydidae and is also one of the most raised species on China’s modern-day turtle farms (da Nóbrega Alves et al., 2008; Jo et al., 2017). Contrastingly, statistically significant greater suitability was observed for the three native *Trachemys*, coinciding with the natural distribution area occupied by each species, again corroborating the much lower establishment risk of TSE regarding the congeners we evaluated.

### Future directions

Our findings show that diverse historical, biogeographical, and ecological processes have defined the current natural and invasive distribution of *T*.*s. elegans*. In this process, its ecological niche has played a key role, showing that in non-native areas it will successfully establish and outcompete a potential competitor if local environmental conditions are closer to its ecological optimum (niche center) than to the ecological optimum of the native species. Moreover, the historical and contemporary patterns of genetic diversity and structure of these closely related *Trachemys* turtles are key to understanding the distribution limits of *T*.*s. elegans*. Our genetic evaluation was enhanced by the novel distribution modeling approach applied, contrasting niche suitability based on niche-center distances between congeners/native and non-native species to explore distribution patterns of their shared suitability space. Finally, our results set the basis for future work, where using whole genome or gene-targeted sequencing and with the inclusion of a higher number of field-sampled individuals, would allow to directly assess hybridization and specific gene introgression and connection between loci and traits.

## Author contributions

SE, EVD, and EMM conceptualized and designed different parts of the research. SE and EVD performed fieldwork and laboratory work. SE, EVD, EAM, IO, and BNR conducted genetic and genomic data analyses. SE, MN, LOO, and EMM performed species niche modeling analyses. SE, EVD, LOO, MN, and FTB wrote the paper, and all authors reviewed the manuscript and agreed on the submission.

## Data accessibility

Sequences (cytochrome *b* and R35) are deposited in GenBank (Accession numbers in Table S1). The microsatellite genotypes and the assembled GBS data (vcf file) used in the study are available on Dryad (or Zenodo) doi: XXX…. Sampling localities, genetic protocols and other data are available in supplementary data.

## Acknowledgments

We are grateful with Juan Fornoni and the members of the Genetics and Ecology laboratory and the Department of Herpetology (AMNH) for helpful discussions along the project. We thank E. Recuero, A. Flores-Manzanero, X. Cortés-Rodríguez, R. Medina, C. Rivera-Arroyo and O. Romero for their help during fieldwork, and R. García Herrera and M. Baltazar for computer systems support. Our gratitude with the Laboratorio de Herpetología FES-Iztacala for facilitating the tissue sampling from living specimens under their care, and the museums AMNH, FMNH and SNM for sharing of the preserved tissue samples. We are in debt with many people along our fieldwork that allowed us to sample on their ejidos and private properties. SEB acknowledges that this paper was a part of his doctoral thesis in the Programa de Doctorado en Ciencias Biológicas de la Universidad Nacional Autónoma de México. SEB had a scholarship and financial support provided by the Consejo Nacional de Ciencia y Tecnología (CONACyT CVU 3646304 / Reg. becario 245447), Program for Postgraduate Studies (PAEP) and UNAM. This project was funded by Consejo Nacional de Ciencia y Tecnología (CONACyT grant # 237228), granted to EVD. The genomic analyses were performed while EVD was on sabbatical at the American Museum of Natural History with financial support from Dirección General de Asuntos del Personal Académico (DGAPA/PASPA 20160609).

## Supplementary data

Supplementary data associated with this article (Appendices S1-S3; Tables S1-S3; Figures S1-S7) can be found, in the online version, at doi: XXX.

