## Supporting data for "Complex genetic patterns and distribution limits mediated by native congeners of the worldwide invasive red-eared slider turtle"

**Supporting Information**
**Appendices S1-S3**
**Tables S1-S3**
**Figures S1-S7**

**Appendix S1**. Extended description of all materials and methods used in the study

### PCR Protocols

#### Microsatellites loci

TSE02-TSE80 primers were amplified as in Xin et al. (2012), in a 5 µl final volume. For Tsc108-330 primers developed by Simison et al. (2013), we designed a multiplex PCR protocol containing ~25ng template DNA, 0.5 units of Taq DNA polymerase, 3mM $MgCl_2$, 0.2 mM dNTPs, 0.45 µM of each primer and 10x reaction PCR buffer (200 mM Tris-HCl pH=8.4, 500 mM KCl). Primer combinations were as follows: 1) Tsc108 & Tsc243; 2) Tsc260 & Tsc328; and 3) Tsc263, Tsc330 & Tsc299.

For all primer pairs, cycling conditions were: initial 5 min denaturation at 95°C; 30 cycles consisting of 94°C for 30 seg, primer-specific annealing temperature for 30 seg and 72°C for 40 seg, and 10 min final extension at 72°C. Annealing temperatures (AT) are in Table S2.

#### Mitochondrial and nuclear sequences

**Cytochrome *b*** (Fritz et al. 2011). The cyt *b* gene was amplified in a 25 µl final volume containing ~25ng template DNA, 0.5 units of Taq DNA polymerase, 2.5mM $MgCl_2$, 0.2 mM dNTPs, 0.4 µM of each primer and 10x reaction PCR buffer (200 mM Tris-HCl pH=8.4, 500 mM KCl). PCR conditions were as follows: initial 5 min denaturation at 94°C; 30 cycles consisting of 94°C for 30 seg, 58°C for 30 seg and 72°C for 1 min, and 10 min final extension at 72°C.

**Genomic intron 1 of the RNA fingerprint protein 35 gene** (Fujita et al. 2004)**.** The R35 gene was amplified in a final volume of 25 µl containing ~25ng template DNA, 0.5 units of Taq DNA polymerase, 3mM $MgCl_2$, 0.2mM dNTPs, 0.5μM of each primer and 10x reaction PCR buffer (200 mM Tris-HCl pH=8.4, 500 mM KCl). PCR conditions were as follows: initial 5 min denaturation at 94°C; 35 cycles consisting of 94°C for 30 seg, 60°C for 1 min 30 seg and 72°C for 2 min, and 10 min final extension at 72°C.

**SNPs calling with *ipyrad***

Considering that the parameters used in *de novo* locus identification and genotyping may affect downstream analyses and resulting inference, we examined a range of values of parameters to optimize assembly in the *ipyrad* pipeline (Eaton and Overcast, 2020); the final processing and filtering protocol used is:

Demultiplex
Assembly method: *denovo*
Restriction overhang (cut1): TGCA
Max low quality base calls (Q<20) in a read: 5
phred Q score offset: 33
Min depth for statistical base calling: 8
Min depth for majority-rule base calling: 8
Max cluster depth within samples: 10000
Clustering threshold for de novo assembly: 0.85
Max number of allowable mismatches in barcodes: 0
Filter for adapters/primers: 1
Min length of reads after adapter trim: 35
Max alleles per site in consensus sequences: 2
Max N's (uncalled bases) in consensus: 5
Max Hs (heterozygotes) in consensus: 8
Min # samples per locus for output: 87
Max # SNPs per locus: 0.2
Max # of indels per locus: 5
Max # heterozygous sites per locus: 0.5

**Phylogenetic analyses**

To obtain a phylogenetic tree incorporating an outgroup, we built a SNP-based outgroup from the Painted turtle *Chrysemys picta* genome (Badenhorst et al. 2015; (ftp://ftp.ncbi.nlm.nih.gov/genomes/all/GCF_000241765.3_Chrysemys_picta_bellii-3.0.3_genomic.fna.gz) with Samtools 1.x (Li 2011).

### Appendix S2. Glossary of mathematical symbols used in niche analysis

| | |
|---|---|
| $A$ | Niche ellipsoid for TSE |
| $\mu, \Sigma$ | Mean vector and covariance matrix for TSE niche |
| $B_k$ | Niche ellipsoid for alternative species $k$ |
| $\mu_k, \Sigma_k$ | Mean vector and covariance matrix for alternative species $k$ |
| $M(p, \mu, \Sigma)$ | Generic Mahalanobis distance between a point $p$ and the mean of multivariate normal distribution with mean $\mu$ and covariance matrix $\Sigma$ |
| $S$ | Generic suitability index defined by $S = e^{-M/2}$, where $M$ is a Mahalanobis distance |
| $S_{1i}$ | Suitability of TSE at the $i$-th presence point of TSE in $A \cap B_k$ |
| $S_{2i}$ | Suitability of species $k$ at the $i$-th presence point of TSE in $A \cap B_k$ |
| $S_{1j}$ | Suitability of TSE at the $j$-th presence point of species $k$ in $A \cap B_k$ |
| $S_{2j}$ | Suitability of species $k$ at the $j$-th presence point of species $k$ in $A \cap B_k$ |
| $f(u, v)$ | Bivariate density function of $(S_{1i}, S_{2i})$ at $(u, v)$ |
| $f_k(u, v)$ | Bivariate density function of $(S_{1i}, S_{2i})$ at $(u, v)$ |
| $D$ | (Quadratic) Distance between densities $f(u, v)$ and $f_k(u, v)$ |
| $\hat{f}(u, v)$ | Kernel density estimate of $f(u, v)$ |
| $\hat{f}_k(u, v)$ | Kernel density estimate of $f_k(u, v)$ |
| $\widehat{D}$ | (Quadratic) Distance between densities $\hat{f}(u, v)$ and $\hat{f}_k(u, v)$ |
| $H_1$ | Area of highest density region (HDR) corresponding to $f(u, v)$ |
| $H_2$ | Area of highest density region (HDR) corresponding to $f_k(u, v)$ |
| $\widehat{H}_1$ | Area of highest density region (HDR) corresponding to $\hat{f}(u, v)$ |
| $\widehat{H}_2$ | Area of highest density region (HDR) corresponding to $\hat{f}_k(u, v)$ |

### Appendix S3. Kernel density estimation and highest density regions

Suppose $(X_1, Y_1), \dots, (X_n, Y_n)$ are sample points of a bivariate random vector, assumed to originate from a two-dimensional density function given by $f(x, y)$. Kernel Density estimation (Scott, 1992; Simonoff, 1996) is a method for inferring the density function, which may be unknown. It is based on the notion of a kernel function, $K$, that is nonnegative and integrates to 1. The *kernel estimate* of $f$ is defined as

$$\hat{f}(x, y) = \frac{1}{n} \sum_{i=1}^{n} \frac{1}{h^2} K\left(\frac{X_i - x}{h}\right) K\left(\frac{Y_i - y}{h}\right),$$

where $h$ is a positive constant called the *bandwidth* that controls the degree of smoothing of the ensuing estimate. A common choice for $K$ is the standard normal density function. Because we are interested in studying bivariate densities restricted to the $(0,1) \times (0,1)$ unit square (what we called the suitability space in the text), a special provision called edge correction must be taken into account. Function `bivariate.density` in R package `sparr` implements kernel density estimation with edge correction to confine estimated density functions to the unit square. In addition, a data-driven bandwidth selection algorithm, that is, a value of $h$ based on data values and an optimality criterion is also provided in that package (Davies et al. 2018).

Next consider an arbitrary bivariate density function, $g(x, y)$. The *highest density region* (HDR) is a useful device for summarizing and describing $g$. It is a subset of the plane that represents location of points of high density spanning a specified probability and is particularly useful for dealing with uncharacteristic and multimodal densities. The $100 \times (1 - \alpha)\%$ HDR (Hyndman, 1996) is the subset $R(c) = \{(x, y) | g(x, y) \geq c\}$, where $c$ is the largest constant such that

$$\iint_{R(c)} f(x, y) \, dx \, dy \ \geq 1 - \alpha.$$

If $g(x, y, z)$ is a three-dimensional density, the HDR is likewise defined to be the subset $R(c) = \{(x, y, z\} | g(x, y, z) \geq c\}$, where $c$ is the largest constant such that

$$\iiint_{R(c)} f(x, y, z)\, dx\, dy\, dz \geq 1 - \alpha.$$

In any case, the HDR is a small set that encompasses high probability. The HDR is a neighborhood of the mode (or modes) of the density having a specific area in two dimensions, or a volume in three dimensions. R library ks provides convenient functions for plotting boundaries of HDRs of given $\alpha$ in two dimensions, as opposed to plotting multiple contour levels of a kernel density estimate, which is a different graphical concept. Since any kernel density estimate is a legitimate density function, then the HDR notion can be directly applied to the estimate as well, as was done in our analysis.

### Tables

**Table S1.** List of *Trachemys* samples used in the study, from fieldwork individuals and from museum specimens, including all distribution and field data, is available as a separate excel file in the Supporting information documents.

**Table S2**. List of microsatellites primers used for the genetic analyses with *Trachemys* species, with their corresponding sequence, annealing temperature applied and source.

| Locus | Primer sequence (5’-3’) | AT (°C) | Published source |
|---|---|---|---|
| **TSE02** | F: TCAGACGTGGCCTTCCTC<br>R: AATCAAACGCTGCTCCCT | 66 | Xin et al. 2012 |
| **TSE06** | F: ACCCTGACATCTGCCGACA<br>R: GAGACCTTCCGCTGCTGC | 68 | Xin et al. 2012 |
| **TSE09** | F: ACGGAGGACACTGCTTGA<br>R: TTGCTTGGCTAAGGTGGA | 64 | Xin et al. 2012 |
| **TSE10** | F: TTTCAAACACCCCTCCAG<br>R: CACCTAGCACCATTTTCC | 60 | Xin et al. 2012 |
| **TSE14** | F: CTGTCGGTGTCTTGTCCC<br>R: TGAGCCCAGAAGTAGTGATG | 64 | Xin et al. 2012 |
| **TSE21** | F: GGAACCGCAAGGAGGAAA<br>R: GCCATGCAACTGAGCACC | 66 | Xin et al. 2012 |
| **TSE78** | F: AAGGCAGCACAAATGGAG<br>R: ACAGAATGTGGCAGGGAC | 66 | Xin et al. 2012 |
| **TSE80** | F: AGACAGTTGCTTCCTTGA<br>R: CATCCCCTTGCTTTTAGT | 60 | Xin et al. 2012 |
| **Tsc108** | F:CGCAGTCAAAACACCTTCAG<br>R:TTCACCTCCCCAGATCTCAC | 55 | Simison et al. 2013 |
| **Tsc243** | F:GCAAAACCTGGAGATTTTCAA<br>R:TTTCGATGGAAAATGGCTTT | 55 | Simison et al. 2013 |
| **Tsc260** | F:TGCAAATGGAGTTGCAAGA<br>R: TCCATTTGAACCTGGGAGAA | 55 | Simison et al. 2013 |
| **Tsc328** | F:TGGATTGCATTATTAGAAATGGT<br>R:CCCACCAACCACCATAATTC | 55 | Simison et al. 2013 |
| **Tsc263** | F:TGTGCACGGGAGTTGTATG<br>R:TTCTATTTGCCAAAAATTGCAT | 55 | Simison et al. 2013 |
| **Tsc330** | F:TGGCTTATTTTGCAGCCTGA<br>R:CCAACTTTCACTCCCATTGC | 55 | Simison et al. 2013 |
| **Tsc299** | F:CCATGTGCCATCTGTCTACCT<br>R:GATCAAGGGATGAGGGTCAA | 55 | Simison et al. 2013 |
| **cyt *b*** | F: GATTTAAGCCGAGACCTGTG<br>R: TCTTTGGTTTACAAGACCAATGC | 58 | Fritz et al. 2011 |
| **R35** | F: ACGATTCTCGCTGATTCTTGC<br>R: GCAGAAAACTGAATGTCTCAAAGG | 60 | Fujita et al. 2004 |

AT: annealing temperature

**Table S3.** List of GenBank accession numbers of mitochondrial cytochrome *b* (cyt *b*) and nuclear genomic intron 1 of the RNA fingerprint protein 35 (R35) sequences used for phylogenetic analyses.

| Species | cyt *b* | R35 |
|---|---|---|
| ***Chrysemys picta*** | HE590298 | HE590495 |
| ***Malaclemys terrapin*** | HE590304 | |
| ***Trachemys venusta*** | HE590362 | |
| ***Trachemys cataspila*** | HE590363 | |
| ***Trachemys decussata*** | HE590331 | |
| ***Trachemys scripta scripta*** | FJ770618<br>FJ770619<br>HE590356<br>HE590358 | |
| ***Trachemys scripta elegans*** | U81351<br>FJ770617<br>FJ770618<br>FJ770619<br>EU787024<br>HE590356<br>HE590358<br>KM216748 | EU787163<br>FJ770705<br>JN707490<br>JN707528<br>HE590519 |

**Table S4.** Results of the ABBA/BABA test (*D* statistics) (Green et al. 2010; Durand et al. 2011). Three hypotheses plus a null model were tested (H1-H3 and Null), using two randomly chosen individuals per species (to test each hypothesis twice) and *Chrysemys picta* as outgroup. TC: *Trachemys cataspila*; TV: *T. venusta*; TSE: *T. scripta elegans*; TT: *T. taylori*.

| **Test** | **H1(A)** | **H2(B)** | **H3(B)** | ***D*** | ***Z*** | ***D stat*** | ***p*** |
|---|---|---|---|---|---|---|---|
| H1a | TC01 | TV45 | TSE9 | -0.1792 | 20.359 | 0.0089 | 0 |
| H1b | TC11 | TV10 | TSE66 | -0.1963 | 22.757 | 0.0086 | 0 |
| H2a | TT02 | TC01 | TSE9 | 0.2748 | 30.64 | 0.0089 | 0 |
| H2b | TT07 | TC11 | TSE66 | 0.0510 | 5.335 | 0.0096 | <0.001 |
| H3a | TV45 | TT02 | TSE9 | -0.0255 | 2.855 | 0.0090 | 0.0043 |
| H3b | TV10 | TT07 | TSE66 | -0.0497 | 5.727 | 0.0087 | <0.001 |
| Null1 | TV45 | TV10 | TV58 | -0.0132 | 2.241 | 0.0059 | 0.025 |
| Null2 | TC01 | TC11 | TC06 | -0.0032 | 0.506 | 0.0063 | 0.612 |

**Table S5**. Niche overlap (%) and observed $\widehat{D}$ and HDRs between *Trachemys scripta elegans* and native species analyzed in the study. Arrows in *p*-values denote if observed values were significant (single-tail: upper (↑) and lower-tailed (↓)).

| | **Overlap (%)** | $\widehat{D}$ | ***p-value*** | $\widehat{H_1}$ | ***p-value*** | $\widehat{H_2}$ | ***p-value*** |
|---|---|---|---|---|---|---|---|
| TSE — *T. venusta* | 23.1 | 77.041 | 0 ↑ | 0.050 | 0.0048 ↓ | 0.045 | 0.0036 ↓ |
| TSE — *T. cataspila* | 66 | 15.676 | 0.0006 ↑ | 0.126 | 0 ↓ | 0.189 | 0.0378 ↓ |
| TSE —*T. taylori* | 100 | 20.809 | 0.139 | 0.152 | 0.091 | 0.080 | 0.0022 ↓ |
| TSE — *A. marmorata* | 95 | 0.888 | 0.0272 ↓ | 0.628 | 0 ↑ | 0.575 | 0 ↓ |
| TSE — *M. reevesii* | 45 | 9.870 | 0 ↑ | 0.292 | 0 ↓ | 0.243 | 0 ↓ |
| TSE — *M. leprosa* | 100 | 4.925 | 0 ↑ | 0.432 | 0 ↓ | 0.506 | 0 ↓ |
| TSE — *E. orbicularis* | 98.5 | 3.034 | 0 ↑ | 0.404 | 0 ↓ | 0.544 | 0 ↓ |
| TSE — *C. longicollis* | 89.6 | 1.594 | 0 ↑ | 0.516 | 0 ↓ | 0.728 | 0 ↑ |
| TSE — *E. macquarii* | 70.5 | 2.186 | 0 ↑ | 0.598 | 0 ↓ | 0.753 | 0.474 |

### Figures

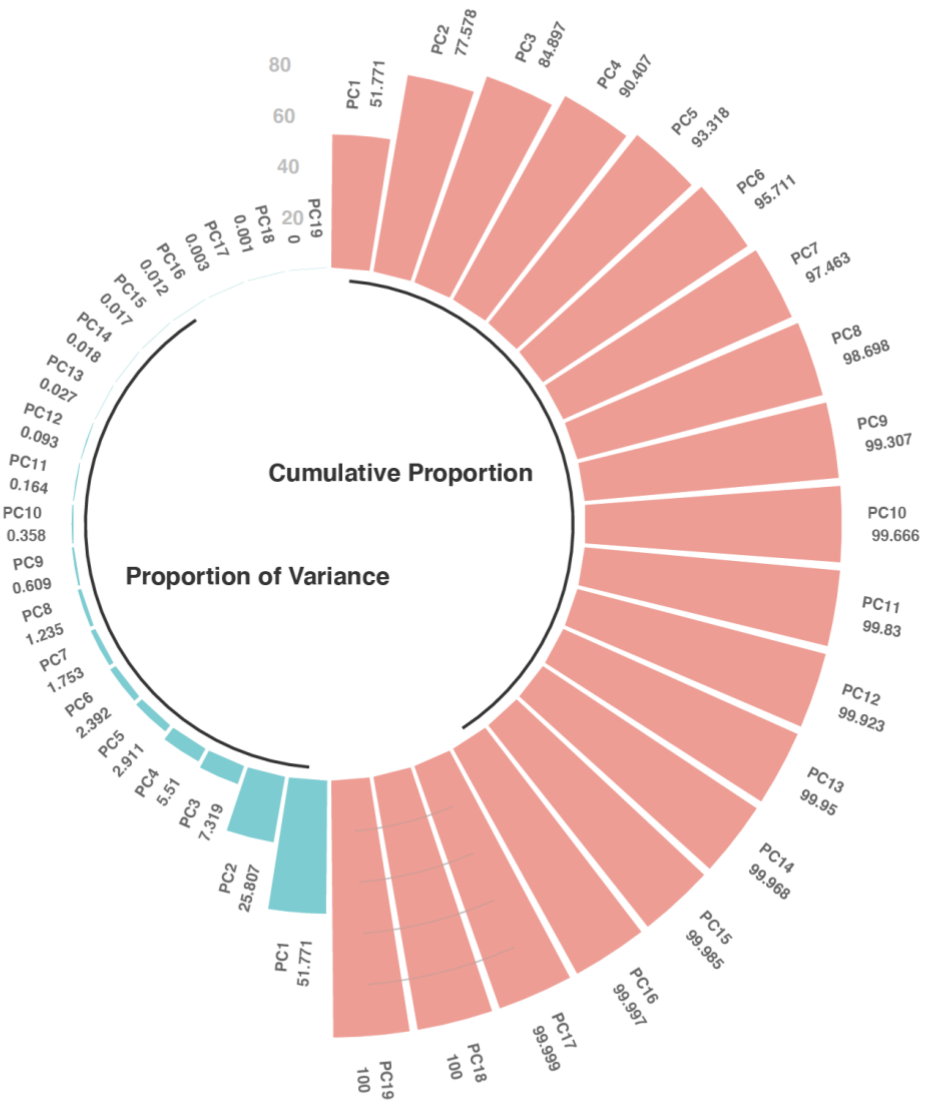


**Figure S1**. Percentage of variance explained by the principal components of the WorldClim 19 bioclimatic variables. Blue bars depict the proportion of variance accounted by each component to the total variance in all the variables. Red bars show the cumulative proportion of variance explained by the first 19 components.

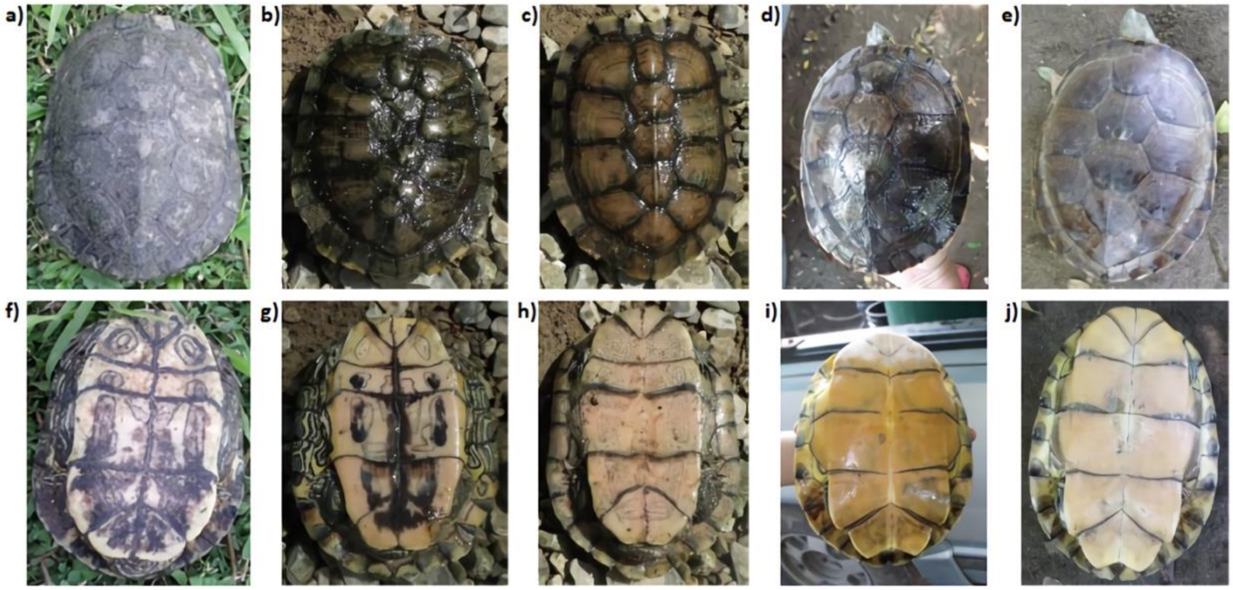


**Figure S2.** Photos of *Trachemys* individuals that could not be identified on the field given their mixed or unusual morphology (from left to right: TSX01, TSX04, TSX05, TSX02, TSX03; Table S1). Variation in color, patterns, and shapes in both dorsal and ventral parts of each turtle is shown. Genomic data allowed to completely differentiate between *T. scripta elegans* (shell: **a-c**; plastron: **f-h**) and *T. venusta* (shell: **d-e**; plastron: **i-j**).

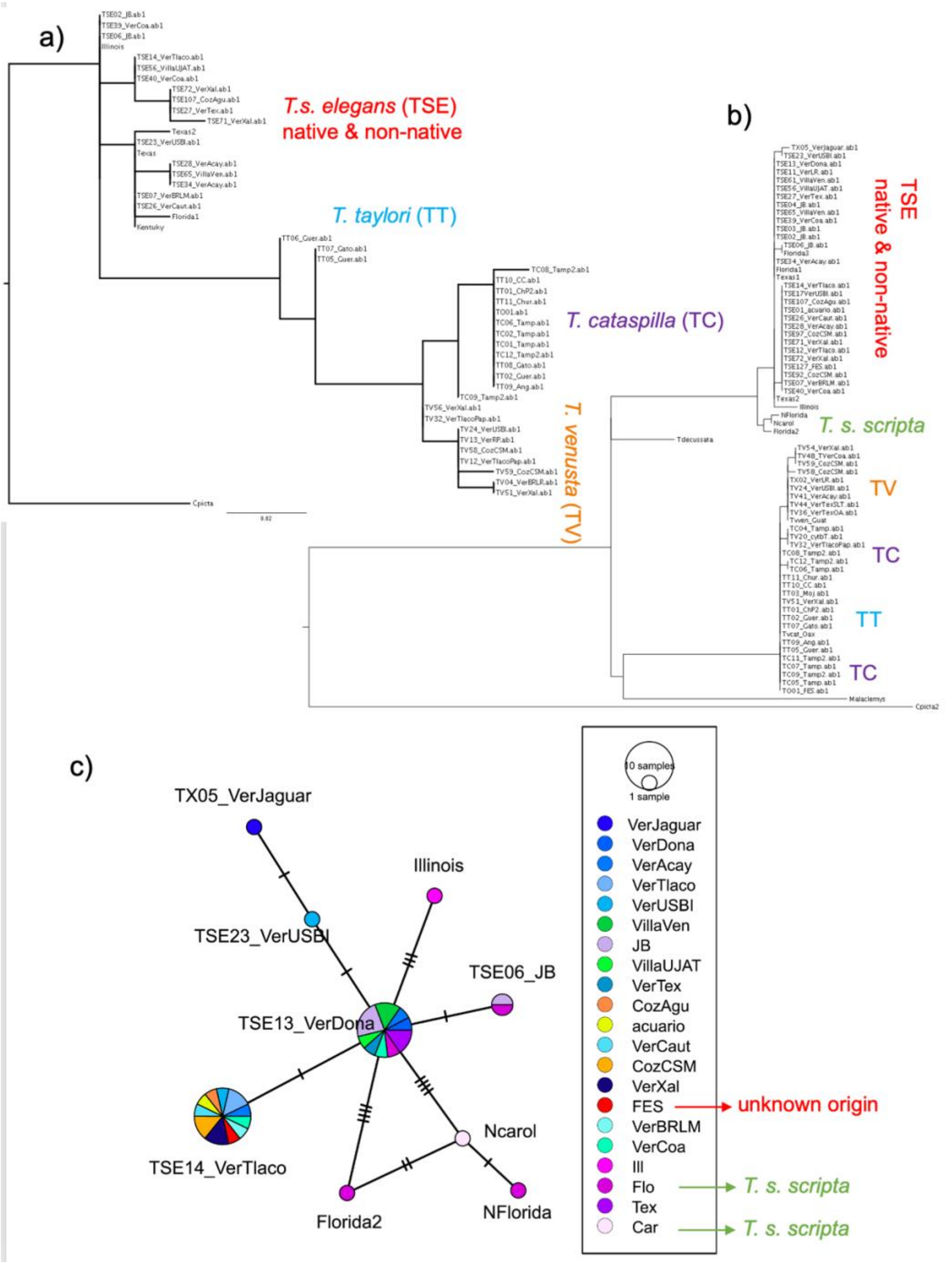


**Figure S3**. Maximum likelihood phylogenetic tree of *Trachemys* species (*T. scripta elegans*, *T. cataspila*, *T. venusta*, *T. taylori*), based on the (**a**) R35 and (**b**) cytochrome *b* genes. *Chrysemys picta* and *Malaclemys terrapin* were used as outgroups. The scale bar represents substitutions per site. (**c**) Minimum spanning haplotype network for *Trachemys scripta elegans* and *T. scripta scripta*. Circles represent haplotypes and circle size is proportional to haplotype frequency. Color of circles depicts the sampling locality or site of origin of samples (see insert; unknown origin, see Table S1).

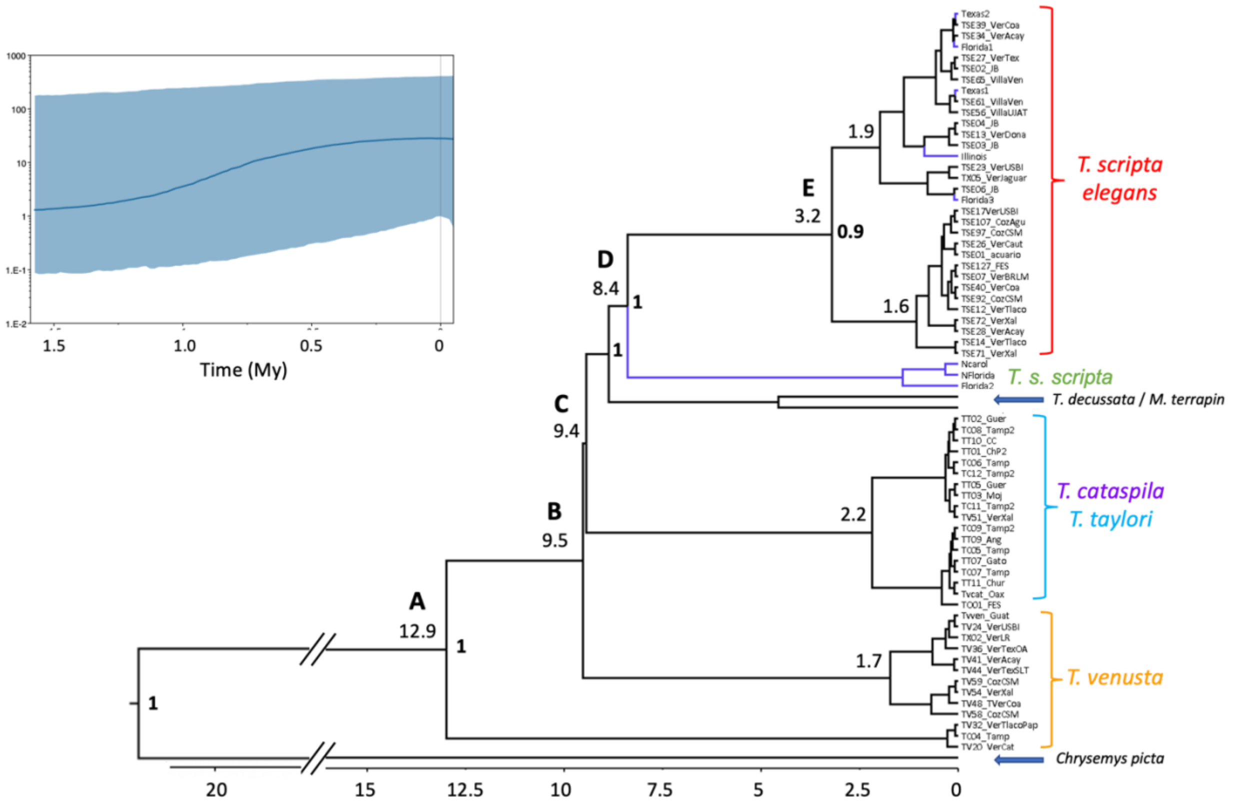


**Figure S4**. Divergence-time estimation (timescale in millions of years; My) above nodes, based on the cytochrome *b* data, of the *Trachemys* species. *Chrysemys picta*, *Malaclemys terrapin* and *Trachemys decussata* sequences were used as outgroups and for calibration points. Posterior probabilities >0.8 are indicated for some nodes, while capital letters are for reference in the results. Skyline plot for non-native *T. scripta elegans* in upper left corner.

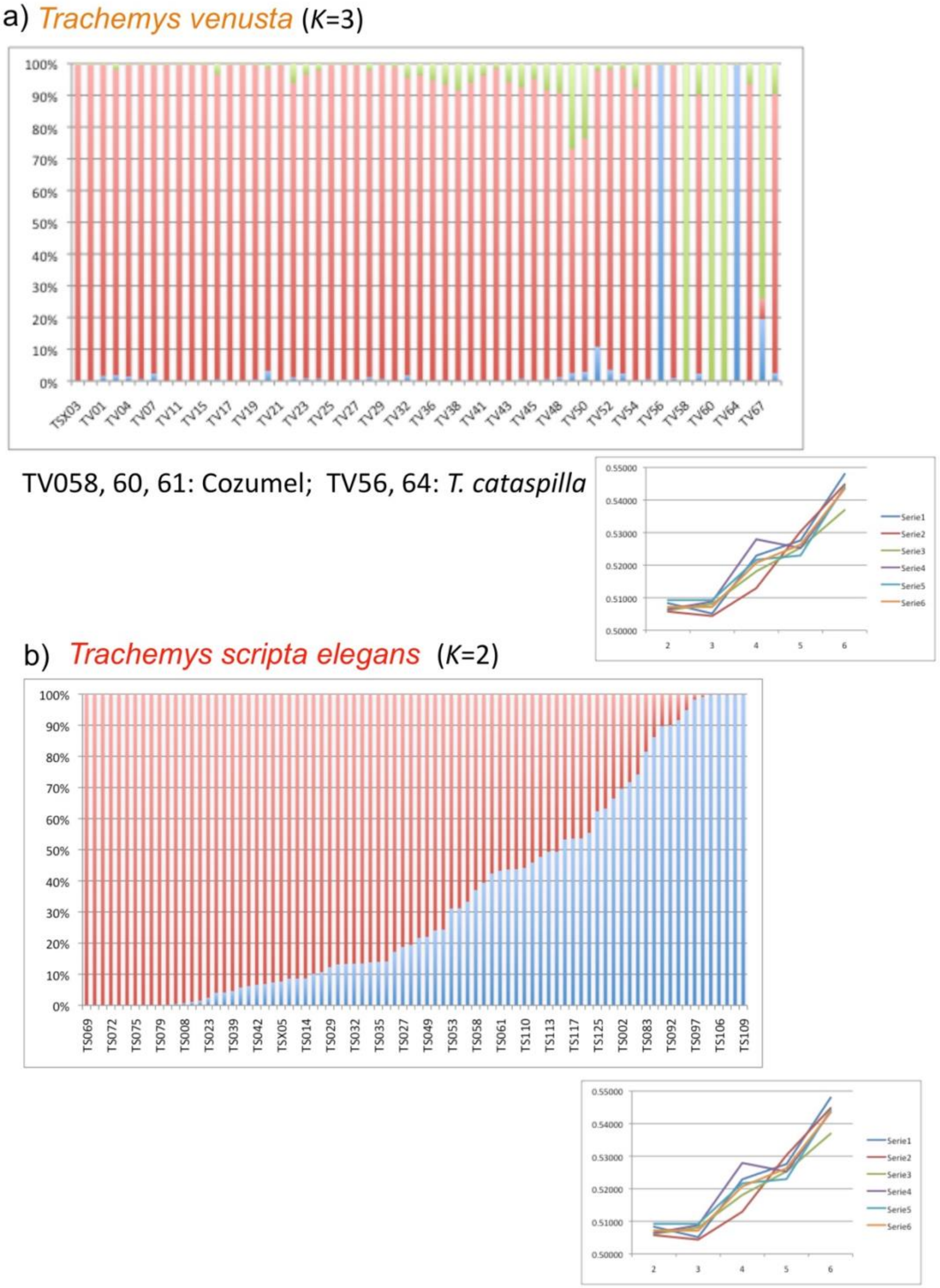


**Figure S5**. sNMF results for (**a**) *Trachemys venusta* (*K* = 3) and (**b**) *T. scripta elegans* (*K* = 2).

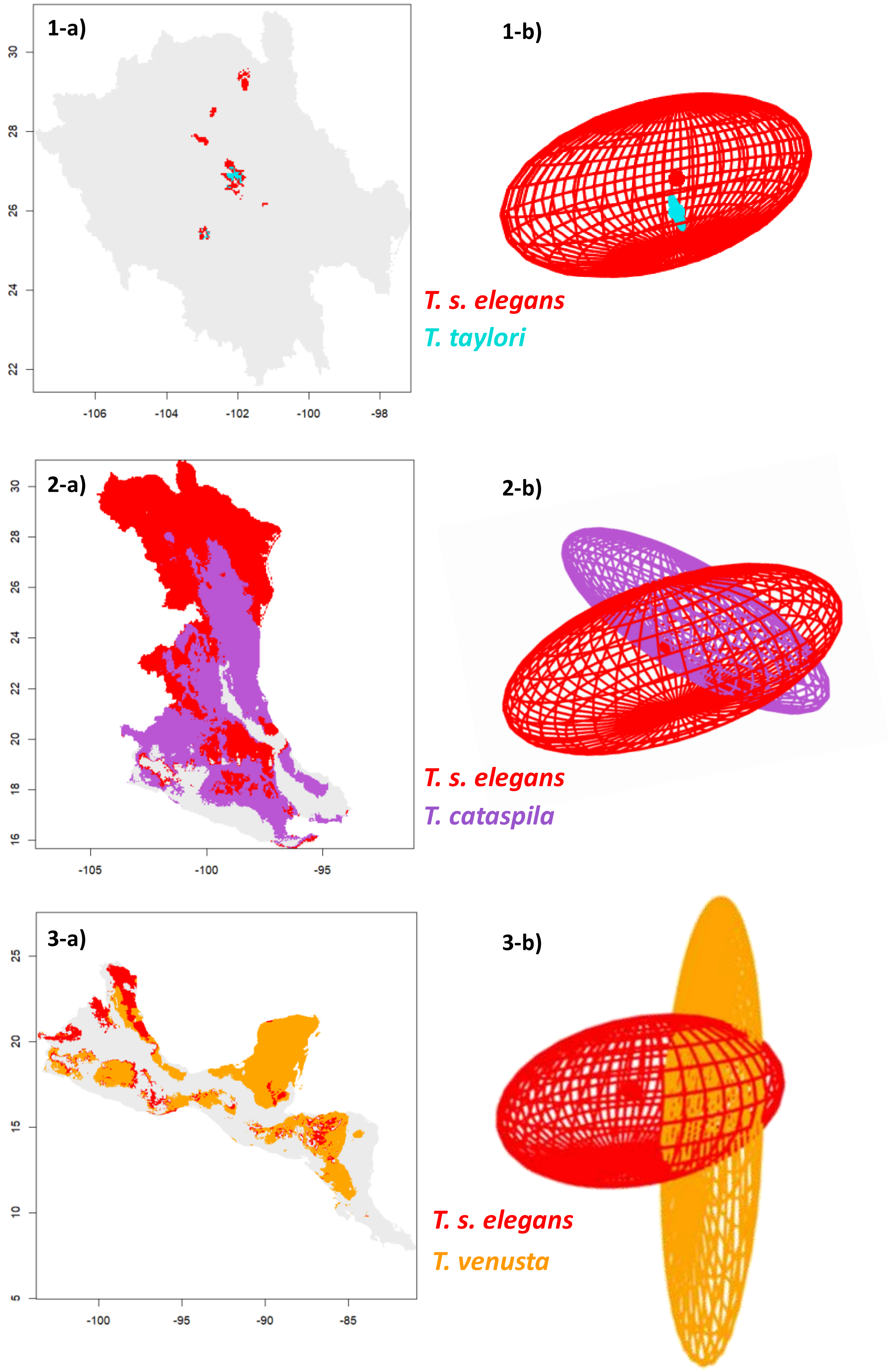
1-a)
30
28
26
24
22
-106
-104
-102
-100
-98
1-b)
T. s. elegans
T. taylori
2-a)
30
28
26
24
22
20
18
16
-105
-100
-95
2-b)
T. s. elegans
T. cataspila
3-a)
25
20
15
10
5
-100
-95
-90
-85
3-b)
T. s. elegans
T. venusta

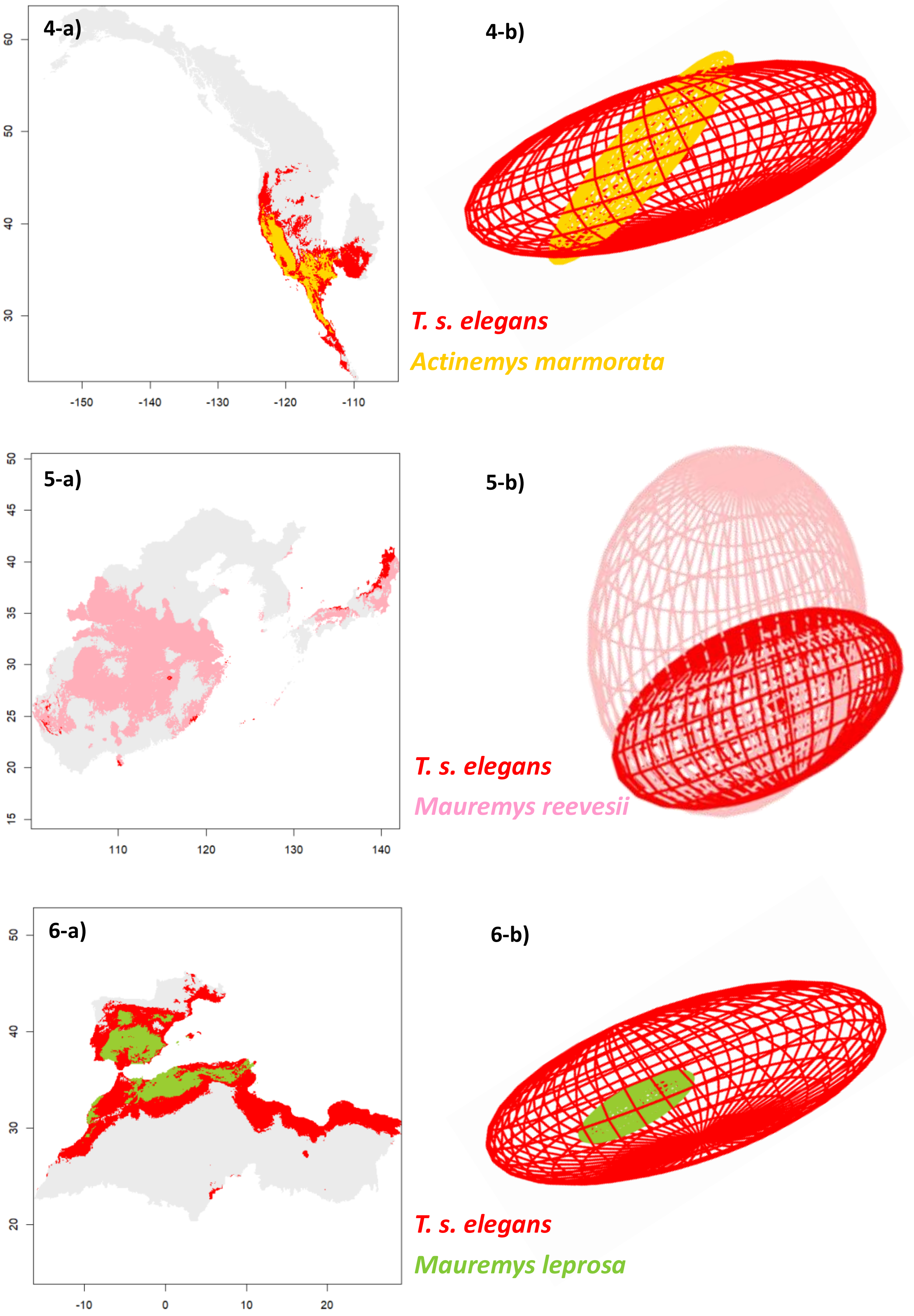
4-a)
4-b)
T. s. elegans
Actinemys marmorata
60
50
40
30
-150
-140
-130
-120
-110
5-a)
5-b)
T. s. elegans
Mauremys reevesii
50
45
40
35
30
25
20
15
110
120
130
140
6-a)
6-b)
T. s. elegans
Mauremys leprosa
50
40
30
20
-10
0
10
20

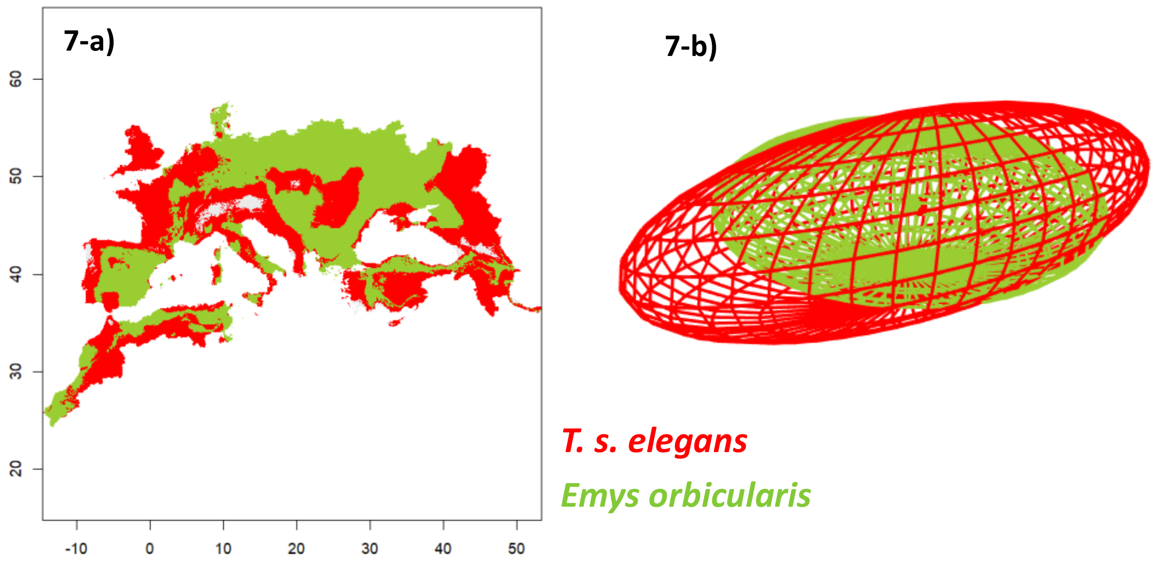
7-a)
60
50
40
30
20
-10
0
10
20
30
40
50
7-b)
T. s. elegans
Emys orbicularis

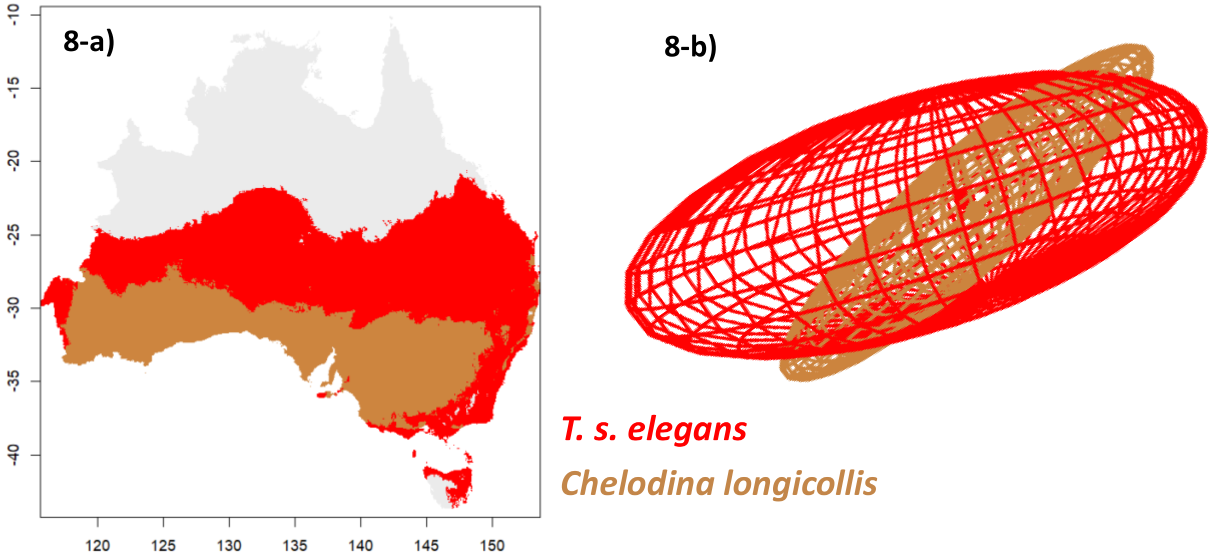
8-a)
-10
-15
-20
-25
-30
-35
-40
120
125
130
135
140
145
150
8-b)
T. s. elegans
Chelodina longicollis

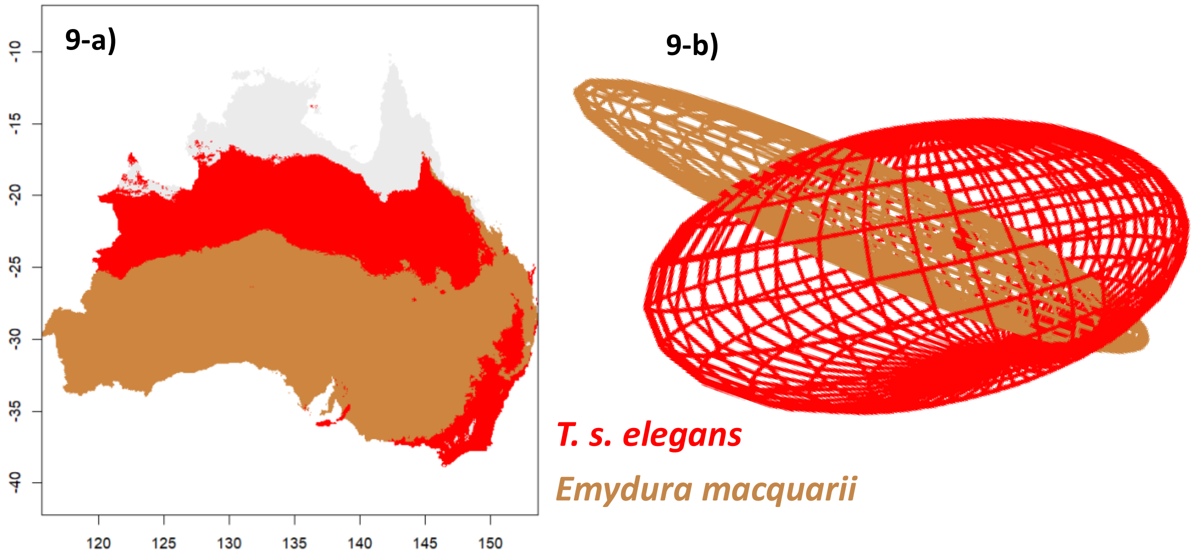
9-a)
-10
-15
-20
-25
-30
-35
-40
120
125
130
135
140
145
150
9-b)
T. s. elegans
Emydura macquarii

**Figure S6.** On the right: Graphic representation of ecological niche overlap, based on ellipsoid models, between *T. scripta elegans* (red) and each native species of different parts of the world: Western United States (4-*Actinemys marmorata*; yellow); Asia (5-*Mauremys reevesii*; pink); Europe (6-*Mauremys leprosa* and 7-*Emys orbicularis*; green), and Australia (8-*Chelodina longicollis* and 9-*Emydura macquarii*; brown). On the left: Maps showing the sites along the distribution of each native species that are closer to the niche-center of *T. scripta elegans*. If the site has a greater suitability (understood as the proximity to the niche-center) for *T. scripta elegans* is colored red; otherwise, it is colored according to each native species color (as mentioned above). Gray color depicts areas where no overlap occurs.

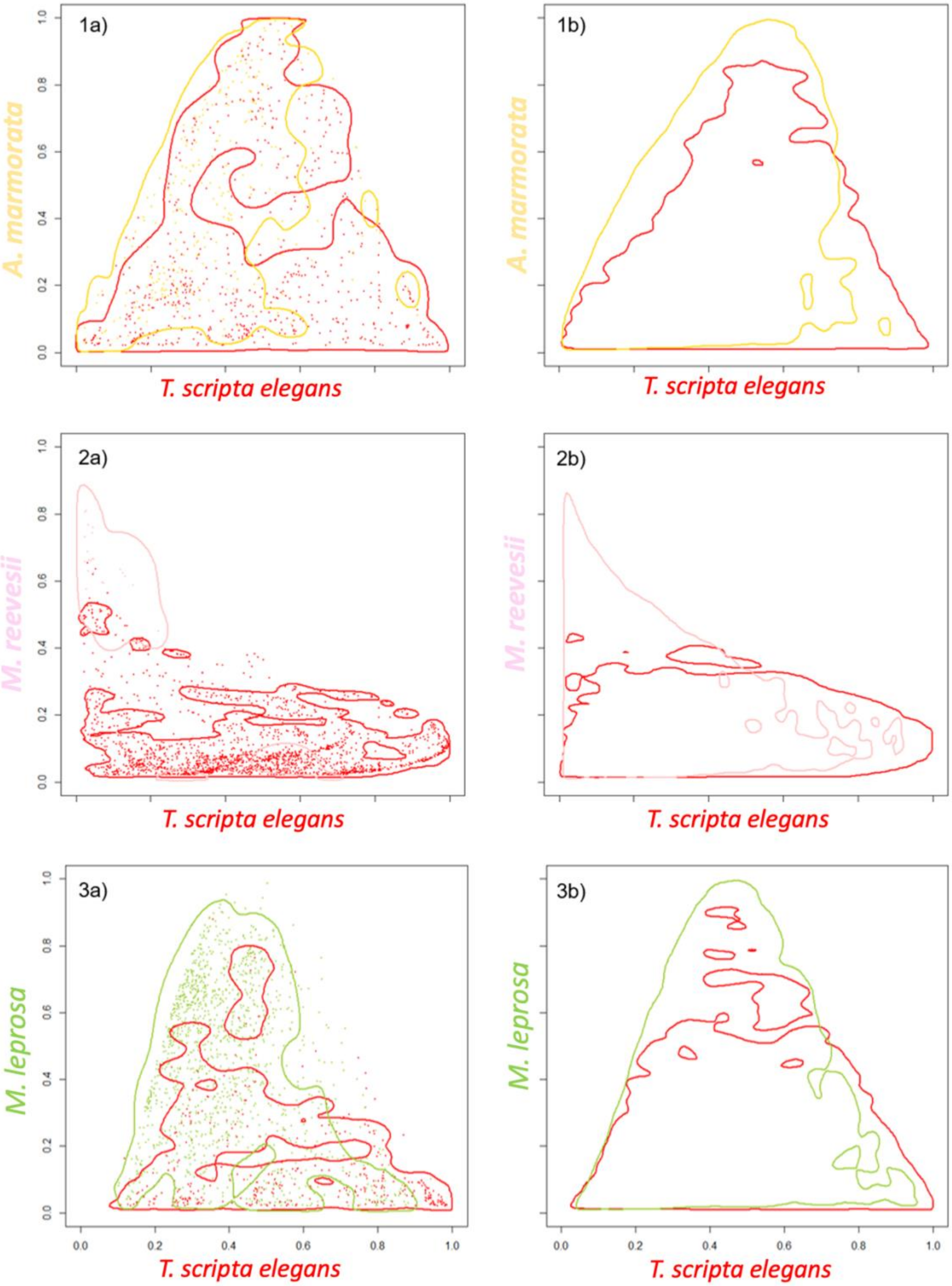

1a)
1b)
2a)
2b)
3a)
3b)
A. marmorata
M. reevesii
M. leprosa
T. scripta elegans
0.0
0.2
0.4
0.6
0.8
1.0

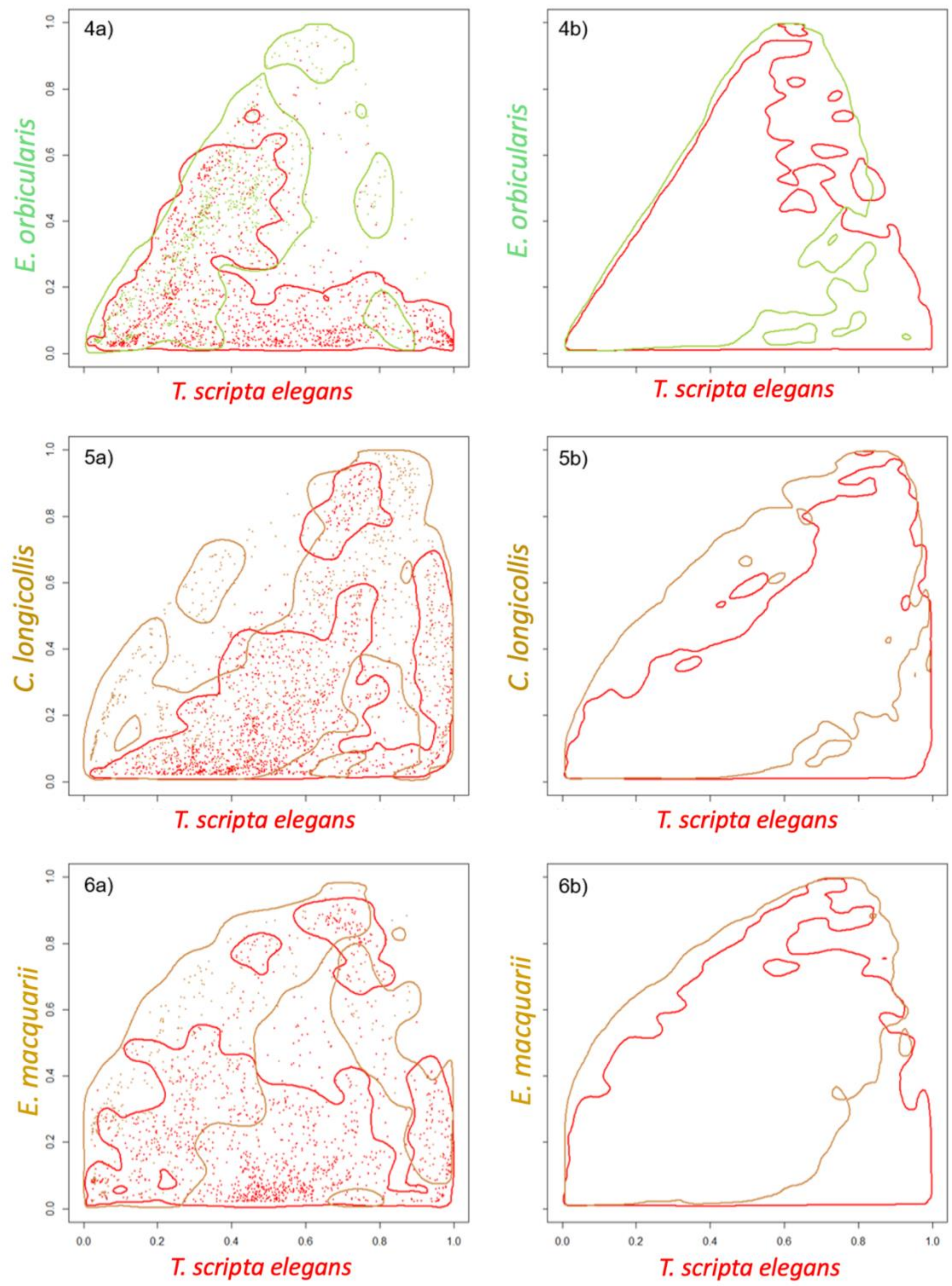


**Figure S7**. Suitability maps for each $k$. The panel on the right shows what the HDRs would be like for densities $f$ and $f_k$ if species truly distributed independently of each other. This panel is obtained by Monte Carlo simulation, generating a large number of independent samples, and applying kernel density estimates. In contrast, the left panel is the observed suitability map based on observed data over $A \cap B_k$.
